# Multi-scale mitochondrial cristae remodeling links Opa1 downregulation to reduced OXPHOS capacity in aged hearts

**DOI:** 10.1101/2025.04.01.644555

**Authors:** Isidora Molina-Riquelme, Gonzalo Barrientos, Leonhard Breitsprecher, Wileidy Gómez, Francisco Díaz-Castro, Silke Morris, Andrea del Campo, Luis Garrido-Olivares, Hugo Verdejo, Olympia Ekaterini Psathaki, Karin B. Busch, Verónica Eisner

## Abstract

Aging is closely associated with cardiovascular diseases, the leading cause of mortality worldwide. Mitochondrial dysfunction is a hallmark of cardiovascular aging because it generates most of the heart’s ATP at the cristae, specialized sub-compartments where OXPHOS takes place. In this study, we used multiple-scale electron microscopy approaches to evaluate age-related mitochondrial and cristae ultrastructural alterations in human and mouse hearts. We found that aged patients’ hearts displayed reduced cristae density as seen by TEM, even before any significant decline in the expression of cristae-shaping proteins. Similarly, a multi-scale approach that included SBF-SEM and TEM showed that in aged mice’s hearts cristae undergo ultrastructural remodeling processes, resulting in a decrease in cristae density and width. Electron tomography suggests an apparent decline in cristae connectivity, and an increase in fenestration size. These changes were linked to Opa1 downregulation, accompanied by reduced OXPHOS maximal respiration, but unrelated to alterations in the levels of OXPHOS core subunits and ATP synthase assembly. Altogether, this indicates that alterations in cristae structure alone are sufficient to impair oxidative metabolism, which highlights its potential as an early signal of cardiac aging, even before noticeable changes in mitochondrial morphology occur.

## INTRODUCTION

The world’s population is progressively aging due to improvements in healthcare and living conditions along with declining birth rates. By 2030, one in six people will be 60 years old or over (1). This demographic trend presents a significant challenge for governments and global organizations, requiring the development of innovative tools to address the needs of the aging population.

Aging is defined as a multifactorial process characterized by a time-dependent decline in function and activity in organisms (2). This leads to the development of age-related diseases, such as cardiovascular diseases (3, 4), the main cause of mortality worldwide (4). Despite their complex etiology, mitochondrial dysfunction has been widely associated with the onset of these diseases (5–7), being a hallmark of cardiovascular aging (8).

Mitochondria are highly dynamic and versatile organelles. Heart mitochondria occupy one-third of the volume of a normal cardiomyocyte and supply 90% of the ATP required by the heart (5). This energy transformation process, as the oxidative phosphorylation (OXPHOS), takes place preferentially at the cristae, specialized sub-compartments formed from folds of the inner mitochondrial membrane (IMM) (9).

Cristae’s structural integrity is essential to maintain mitochondrial function. Cristae shaping determines the assembly of the OXPHOS machinery into complexes and supercomplexes (supramolecular arrangements of the OXPHOS complexes), ensuring its respiratory efficiency which increases ATP synthesis (10). Additionally, cristae can widen or narrow upon changes in metabolic demands (10–13) and cues in the medium (14) to modulate their bioenergetic activity. Cristae shape also regulates cell survival, as their opening allows for the release of cytochrome c to the cytosol, a proapoptotic event (15).

Cristae shape is determined by the concerted action of three molecular entities: mitochondrial contact site and cristae organizing system (MICOS), OPA1, and ATP synthase (9, 16). The MICOS complex in association with OPA1 stabilizes the curvature at the opening of the cristae, forming the cristae junction (9, 16). ATP synthase forms dimers and oligomers that stabilize the curvature at the tip of the cristae (9, 16). Furthermore, different processes influence the shape of the cristae. For instance, alterations in mitochondrial fusion (17) and fission (18) dynamics, named mitochondrial meso-dynamics, are associated with changes in cristae shape, therefore, determining mitochondrial nano-dynamics.

Defects in cristae morphology are associated with aging across different organisms. In aged *Podospora anserina*, ATP synthase dimers dissociate, inducing the collapse of cristae and the emergence of vesicles (19). In aged *Drosophila melanogaster*, mitochondria showed aberrant phenotypes and widened cristae, while mitochondria from aged mouse liver exhibited spaces devoid of cristae (20). In cardiomyocytes of aged sheep, a loss of cristae integrity was correlated with a decrease in the levels of the proteins Opa1 and Mic60 (21). In rat neonatal ventricular cardiomyocytes (NVCM) and human induced pluripotent stem cells (hiPSC)-derived cardiomyocytes, we demonstrated that the induction of cellular senescence using low doses of doxorubicin produces cristae alterations, which was correlated with a decrease of the OXPHOS function and membrane potential (22). In aged mouse heart, a decrease in cristae number (23, 24), aberrant cristae folding, and changes in the cristae tip curvature (23) have been described. Whether these changes are also visible in the human heart is currently unknown, although cristae shape alterations are present in several myopathies (25).

In this work, we aimed to investigate whether changes in cristae shape were observed in a naturally aged model. To study this, we compared the mitochondrial ultrastructure of the atria of human adult (<65 years) and aged (≥65 years) male patients, and the ventricle of adult (5-6 months) and aged (17-19 months) male mice. We applied a multi-scale approach, including transmission electron microscopy (TEM), serial block face-scanning electron microscopy (SBF-SEM), and electron tomography, revealing a consistent pattern between human and mouse mitochondria. Namely, we found a decrease in cristae density with aging. Those changes were independent of cristae shaping proteins in humans but were related to diminished expression of Opa1 in mice, with a reduction in OXPHOS maximal respiration. These findings suggest that alteration in cristae shape impacts negatively in OXPHOS capacity, and precedes noticeable changes in mitochondrial morphology, as an early hallmark of aging in the heart.

## RESULTS

### Mitochondrial and cristae ultrastructural remodeling in aged human hearts

Aging is strongly linked to mitochondrial dysfunction, resulting in decreased ATP output and increased ROS production (5). Given that cristae are crucial for both of these processes (10), we investigated mitochondrial and cristae ultrastructure alterations in the hearts of aged human individuals. To address this question, we collected right atrial appendage tissue from male patients and defined 65 years old as the threshold to distinguish between adults and aged patients. The two subsets of individuals showed comparable clinical metrics, uncovering the absence of a striking cardiac phenotype in the aged individuals (**Table 1**).

**Table 1.** Clinical characteristics of patients in the study.

|  | **Adult** | **Aged** | **P-value** |
| --- | --- | --- | --- |
| **Number of patients** | 5 | 7 |  |
| **Age (years)** | 56 ± 3<br>(49 - 64) | 72 ± 2<br>(65 - 78) | 0.0004 |
| **Height (m)** | 1.71 ± 0.02<br>(1.65 - 1.77) | 1.71 ± 0.03<br>(1.64 - 1.86) | 0.9785 |
| **Weight (kg)** | 89 ± 9<br>(57 - 108) | 78 ± 5<br>(65 - 103) | 0.3021 |
| **BMI (kg/m<sup>2</sup>)** | 30.1 ± 2.8<br>(20.9 - 36.5) | 26.6 ± 1<br>(23.2 - 29.8) | 0.2052 |
| **LVEF (%)** | 58 ± 3<br>(50 - 65) | 56 ± 2<br>(50 - 60) | 0.5404 |
| **LV thickness** | 12 ± 1<br>(10 - 13) | 11 ± 1<br>(10 - 14) | 0.9999 |
| **Septum thickness** | 13 ± 1<br>(11 - 15) | 13 ± 1<br>(10 - 16) | 0.7524 |
| **LA diameter (mm)** | 43 ± 4<br>(36 - 53) | 43 ± 1<br>(39 - 45) | 0.9470 |
| **Preoperative creatinine (mg/dL)** | 1.11 ± 0.13<br>(0.8 - 1.53) | 1.12 ± 0.07<br>(0.9 - 1.46) | 0.9489 |
| **Extracorporeal circulation (min)** | 150 ± 4<br>(141 - 164) | 150 ± 8<br>(117 - 185) | 0.9958 |
| **Aortic clamp (min)** | 106 ± 5<br>(94 - 116) | 109 ± 9<br>(88 - 150) | 0.8430 |
| **Arterial hypertension (%)** | 80 | 86 |  |
| **Atrial fibrillation (%)** | 0 | 0 |  |
| **History of acute MI (%)** | 0 | 57 |  |
| **Diabetes mellitus (%)** | 40 | 43 |  |
| **Hypercholesterolemia (%)** | 80 | 57 |  |
| **Beta blockers (%)** | 40 | 71 |  |

Data are presented as mean ± SE or %. The range is presented in parentheses. Data following a normal distribution (P<0.05 with Shapiro-Wilk test) was analyzed with a t-Student test to determine statistical significance. Otherwise, a Mann-Whitney test was used. Data from at least 4 patients for LVEF, LV and septum thickness and LA diameter.

We used 2D TEM to study mitochondrial ultrastructure (**Figure 1A**, compare arrowheads). Our morphometric analysis showed unaltered mitochondrial area in the aged individuals (**Figure 1C**), with an increase in roundness (**Figure 1D**), indicating that mitochondrial shape in cardiomyocytes is lost in aging. Mitochondrial morphology alterations are associated with several pathologies, as mitochondrial diseases, bacterial infection, cancer or cell death, and are commonly associated to imbalance fusion and fission events (26). Particularly, fusion is determined by the GTPases MFN1, MFN2, and OPA1, while fission by the proteins DRP1, MFF, FIS1, MID49, and MID51 (27). Interestingly, the change in ultrastructure was not explained by alterations in the mitochondrial meso-dynamics machinery, since we did not observe changes in the transcript nor the protein levels of MFN1 and OPA1, or in the transcript levels of *DRP1* and *MFF* (**Figure 1, G-I**). However, the lack of statistical differences in mRNA and protein levels may be due to the intrinsic variability of the human samples.

**Figure 1.**
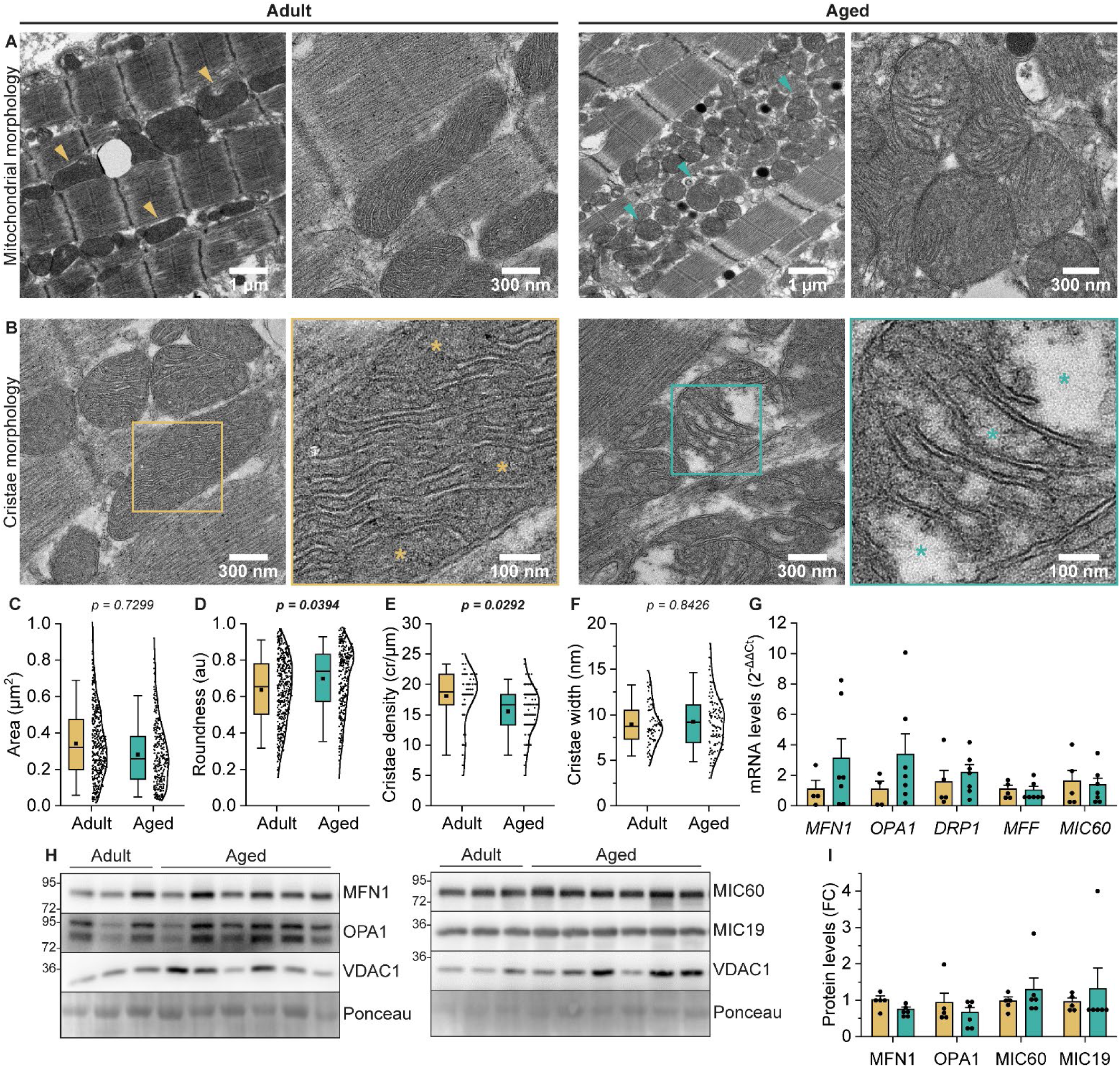
2D TEM shows that mitochondrial and cristae ultrastructure is altered in aged human hearts. **(A-B)** Representative TEM images of mitochondrial **(A)** and cristae **(B)** morphology from right atrial appendage samples obtained from human patients. Adult patients were classified as <65 years old, while aged patients were classified as ≥65 years old. Arrowheads highlight mitochondrial morphology. Asterisks highlight regions devoid of cristae. **(C)** Mitochondrial area (µm^2^). **(D)** Mitochondrial roundness (au). **(E)** Cristae density (cristae/µm). **(F)** Cristae width (nm). **(G)** mRNA levels expressed as 2^−ΔΔCt^ against r18S. **(H)** Western blot of mitochondrial fusion (left) and cristae shaping (right) proteins. **(I)** Protein levels from **(H)** expressed as fold change (FC), normalized against Ponceau staining as a loading control. For TEM analysis, samples from 2 adult and ≥3 aged patients were analyzed. The number of mitochondria analyzed was: Area: 421 and 289; Roundness: 424 and 296; Cristae density: 80 and 120; Cristae width: 80 and 120. Data not following a normal distribution (P<0.05 with Shapiro-Wilk test) was normalized using the rankit procedure. A hierarchical mixed random intercept model was used according to (73) to determine statistical significance. The details of the hierarchical data distribution are presented in **Supplemental Figure 1A**. For RT-qPCR and WB, samples from ≥4 adult and 7 aged patients were analyzed. Data following a normal distribution (P>0.05 with Shapiro-Wilk test) was analyzed with a t-Student test to determine statistical significance. Otherwise, a Mann-Whitney test was used. For box plots: box: P25 and P75; whiskers: P5 and P95; line: median; square: mean. Every point represents a single mitochondrion. For bar plots: mean ± SEM. Each point represents a patient.

Next, we aimed to study cristae ultrastructure (**Figure 1B**, compare asterisks). We found a reduction in cristae density in the aged individuals (18.06 ± 0.49 nm in adult and 15.52 ± 0.39 nm in aged individuals, P=0.0292), indicating a concomitant reduction in the surface of the compartment in which OXPHOS can occur (**Figure 1E**), while cristae width did not show differences between groups (**Figure 1F**). Finally, the transcript levels of cristae-shaping proteins were unaltered, including *OPA1* and *MIC60* (**Figure 1G**), consistent with the protein levels of OPA1, MIC60, and MIC19 (**Figure 1, H and I**).

Functional data from the aged participants in this study showed that those patients did not exhibit significant cardiac dysfunction (**Table 1**). However, as seen by electron microscopy, mitochondria displayed altered morphology and diminished cristae density, linked to a mechanism independent of mitochondrial dynamics and cristae shaping protein levels.

### Mitochondrial morphology is robust in aged mice hearts

We previously demonstrated that alterations in mitochondrial morphology are a common feature in different models of cardiac senescence, such as rat NVCM and hiPSC-derived cardiomyocytes (22). To evaluate if these alterations were also observed in a mouse model of aging, we examined the apex of the left ventricle, harvested from 6 months (adult) and 17-19 months (aged) C57Bl6/J male mice using various electron microscopy techniques. Aged mice exhibited a significant increase in body weight when compared to the adult group (**Table 2**). The aged group also presented a significantly increase in the cardiac hypertrophy index, calculated as the heart weight and tibia length ratio (**Table 2**), indicating that these mice showed this classic cardiac aging hallmark (28).

**Table 2.**
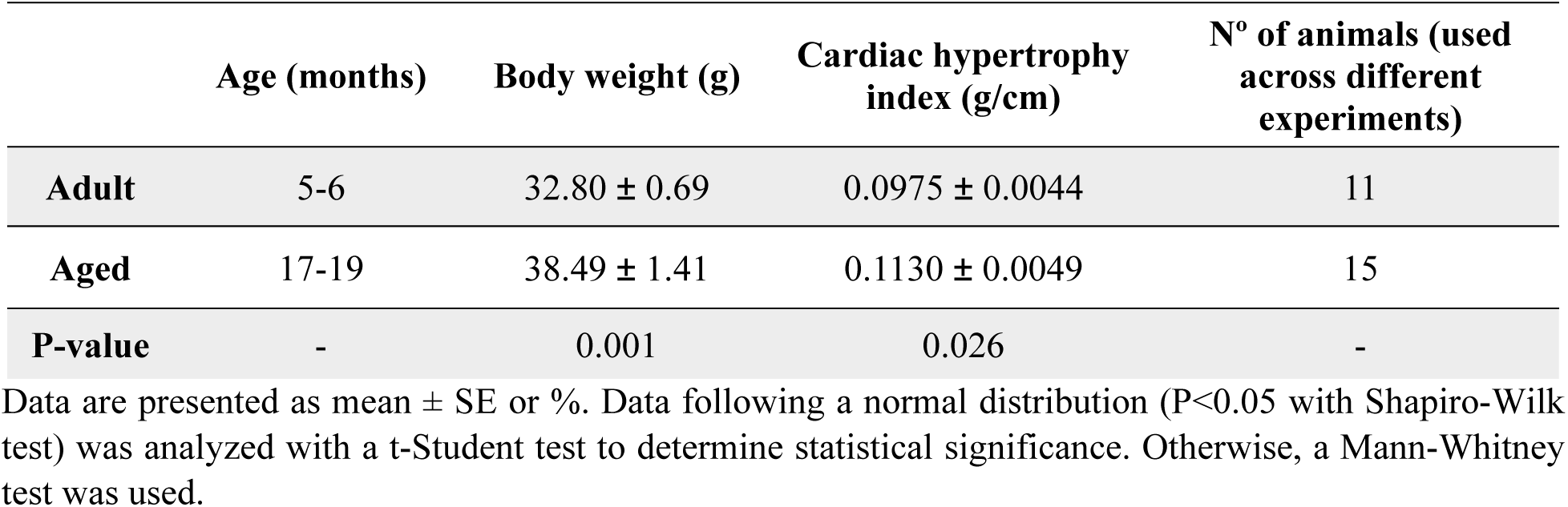
Mice characterization on physical parameters and cardiac hypertrophy.

|  | Age (months) | Body weight (g) | Cardiac hypertrophy index (g/cm) | N° of animals (used across different experiments) |
| --- | --- | --- | --- | --- |
| **Adult** | 5-6 | 32.80 ± 0.69 | 0.0975 ± 0.0044 | 11 |
| **Aged** | 17-19 | 38.49 ± 1.41 | 0.1130 ± 0.0049 | 15 |
| **P-value** | - | 0.001 | 0.026 | - |

Data are presented as mean ± SE or %. Data following a normal distribution (P<0.05 with Shapiro-Wilk test) was analyzed with a t-Student test to determine statistical significance. Otherwise, a Mann-Whitney test was used.

First, we analyzed our samples using 2D TEM (**Figure 2A**, compare arrowheads), and detected a parallel distribution of mitochondria along the contraction axis, according to the classical morphology observed in cardiac muscle (29). We calculated the mitochondrial area (µm^2^) (**Figure 2B**) and roundness (**Figure 2C**), neither of which showed statistical differences between conditions. These results suggest that mitochondrial morphological changes observed in aged human tissue do not exactly replicate in our murine aging model.

**Figure 2.**
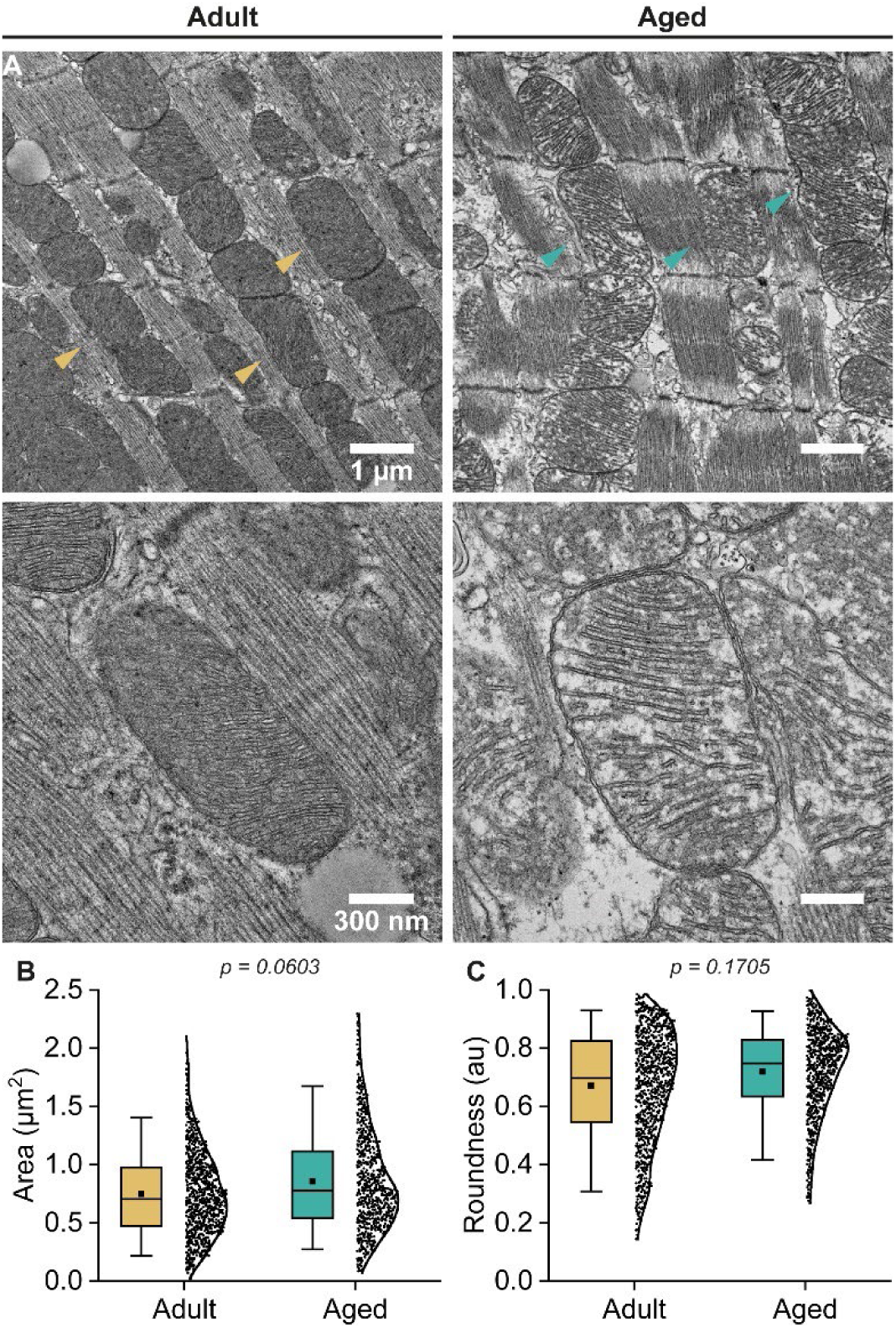
2D TEM shows that mitochondria maintain area and roundness in aged mice hearts. **(A)** Representative images of mitochondrial morphology from adult and aged mice hearts. Arrowheads highlight mitochondrial morphology. **(B)** Mitochondrial area (µm^2^). **(C)** Mitochondrial roundness (au). Samples from 5 animals per condition were analyzed. Data not following a normal distribution (P<0.05 with Shapiro-Wilk test) was normalized using the rankit procedure. A hierarchical mixed random intercept model was used according to (73) to determine statistical significance. The details of the hierarchical data distribution are presented in **Supplemental Figure 1B**. The number of mitochondria analyzed was: Area, 956 and 747; Roundness, 967 and 758. For box plots: box: P25 and P75; whiskers: P5 and P95; line: median; square: mean. Every point represents a single mitochondrion.

Then, we employed SBF-SEM, a high-volume three-dimensional electron microscopy imaging approach that allows the inspection of broad cellular regions, as opposed to 2D TEM field restricted imaging (30). We analyzed tissue samples comprising a volume of 20×20×6 µm^3^, with at least three technical replicates per biological sample. As in 2D TEM analysis, we focused our attention on intermyofibrillar mitochondria (IMF) for two reasons: first, to analyze comparable regions of the ventricular myocytes, and second, because this mitochondrial subpopulation is proposed to be particularly impaired in aging (5, 31). Thus, from every stack of images, we created a sub-volume of 9.5×9.5×6 μm^3^ by cropping the dataset to focus on regions enriched on the IMF population (**Supplemental Figure 2, A and B**). This allowed us to study at least 500 mitochondria per condition and perform different morphological analyses (**Supplemental Figure 2C**). Initial automatic segmentation was performed using Empanada, a Napari plugin for panoptic detection of organelles in electron microscopy images (32), followed by semi-automatic refinement for quality control (**Supplemental Figure 2D and Supplemental Figure 3A, Supplemental Video 1 and 2**, for adult and aged condition, respectively).

Using our segmented model, we analyzed morphometric aspects of the mitochondrial network (**Figure 3A**), finding that adult and aged mice showed no significant difference in volume density (µm^3^/µm^3^) (**Figure 3C**) and surface density (µm^2^/µm^3^) (**Figure 3D**). This indicates that the volume and surface occupied by the mitochondrial network remained similar between conditions. Regarding morphometric aspects of individual IMF, we found no significant differences in individual volume (µm^3^) (**Figure 3E**) and individual surface (µm^2^) (**Figure 3F**) between adult and aged animals.

**Figure 3.**
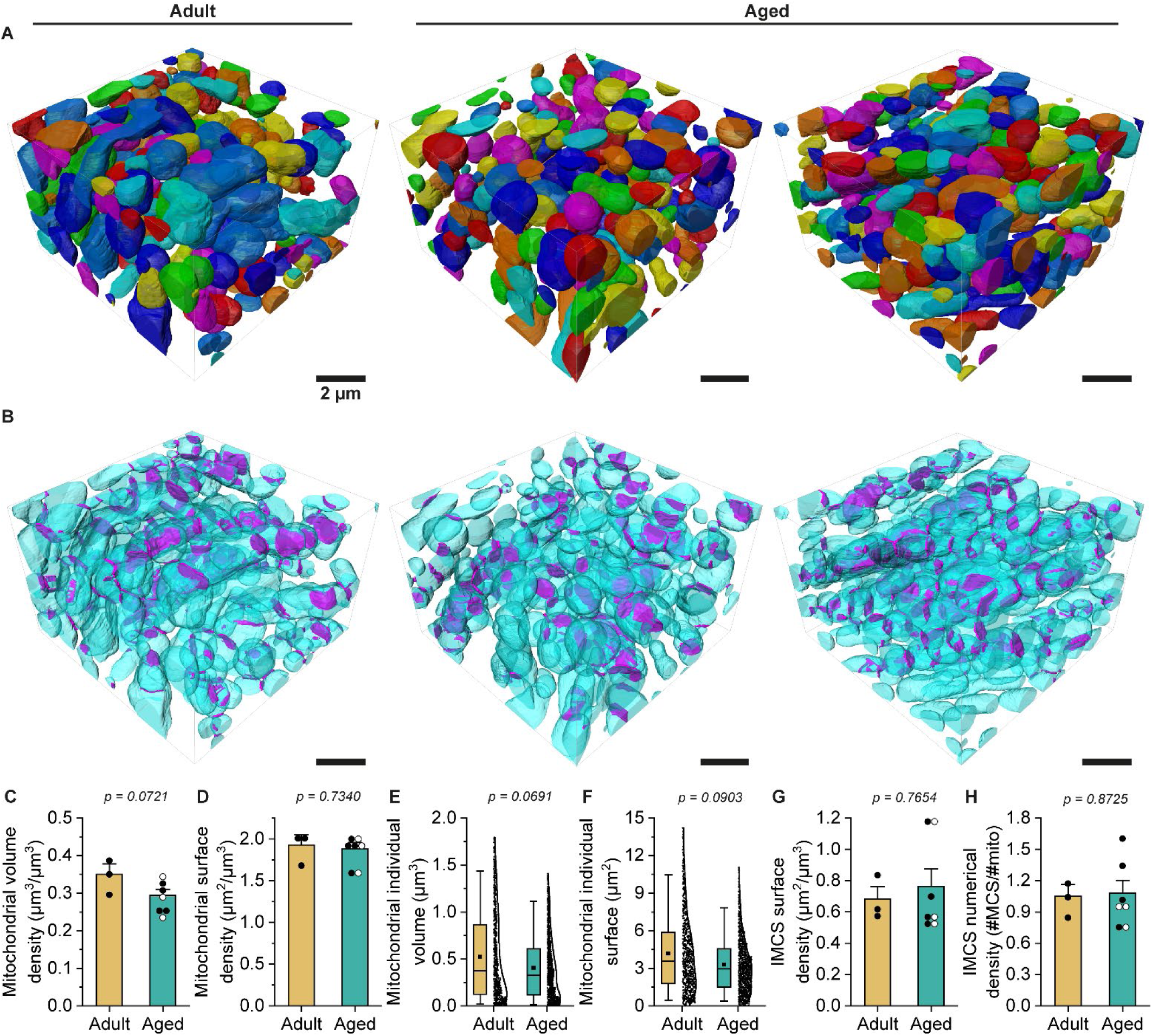
3D SBF-SEM shows that mitochondria maintain density, size, and connectivity in aged mice hearts. **(A-B)** 3D rendering of adult and aged mice heart mitochondria **(A)** and inter mitochondrial contact sites (IMCS) **(B)**. Mitochondria are depicted in multiple colors **(A)** or cyan **(B)**, while IMCS in magenta. The sub-volume is 9.5×9.5×6 µm^3^ with a voxel size of 2×2×25 nm^3^. **(C)** Mitochondrial volume density (µm^3^/µm^3^). **(D)** Mitochondrial surface density (µm^2^/µm^3^). **(E)** Mitochondrial individual volume (µm^3^). **(F)** Mitochondrial individual surface area (µm^2^). **(G)** IMCS surface density (µm^2^/µm^3^), defined as the summation of all mitochondrial contact site area (µm^2^) divided by the summation of all mitochondrial volume (µm^3^) per stack. **(H)** IMCS numerical density, calculated as the number of contact sites per number of mitochondria in each stack. Samples from 1 adult (3 subsamples) and 2 aged (3 and 4 subsamples) animals were analyzed. For **E** and **F**, the number of mitochondria analyzed was: Mitochondrial individual volume, 585 and 1644; Mitochondrial individual surface, 651 and 1775. Data not following a normal distribution (P<0.05 with Shapiro-Wilk test) was normalized using the rankit procedure. A hierarchical mixed random intercept model was used according to (73) to determine statistical significance. The details of the hierarchical data distribution are presented in **Supplemental Figure 1C**. For box plots: box: P25 and P75; whiskers: P5 and P95; line: median; square: mean. Every point represents a single mitochondrion. For bar plots: mean ± SEM. Every point represents a sub-volume (18 and 19 months represented as white and black points, respectively).

Next, using the MCcal-Plugin in MIB (33) we detected inter-mitochondrial contact sites (IMCS) (**Supplemental Figure 2E and Supplemental Figure 3B**), defined here as two or more mitochondria in close vicinity (<100 nm). Based on the segmentation of individual mitochondria we created a map for the IMCS (**Figure 3B**), finding no differences between the adult and the aged mice groups in terms of surface density (µm^2^/µm^3^) (**Figure 3G**) and numerical density (**Figure 3H**). This indicated that mitochondrial network connectivity remained unaltered in the hearts of aged animals.

In sum, our data show that the morphology of the mitochondrial network and individual mitochondria from the murine left ventricle remains unaltered in aged individuals, opposite to our results in human samples. Additionally, we showed that mitochondrial connectivity is conserved, since we did not detect changes in IMCS, pointing to a robust mitochondrial network in aged mice.

### Cristae morphology is altered at a multi-scale level in aged mice hearts

We investigated whether the changes observed at the cristae level in our human samples were also present in mice. For this, we used 2D TEM (**Figure 4A**, compare asterisks and arrowheads), finding differences between the aged and adult conditions. Cristae density was significantly reduced in the aged group; 22.76 ± 0.25 and 21.29 ± 0.20 cristae/um in adult and aged cardiac mitochondria, respectively (P=0.0008) (**Figure 4B**). Strikingly, the cristae width was significantly narrower in the cardiac mitochondria of aged animals (7.40 ± 0.18 nm) as opposed to the adult group (10.23 ± 0.46 nm) (P=0.0372) (**Figure 4C**). These results show a consistent pattern of an overall decrease in the surface of cristae, the IMM-specialized platforms for ATP production, in the mitochondria of aged mice and human cardiac mitochondria.

**Figure 4.**
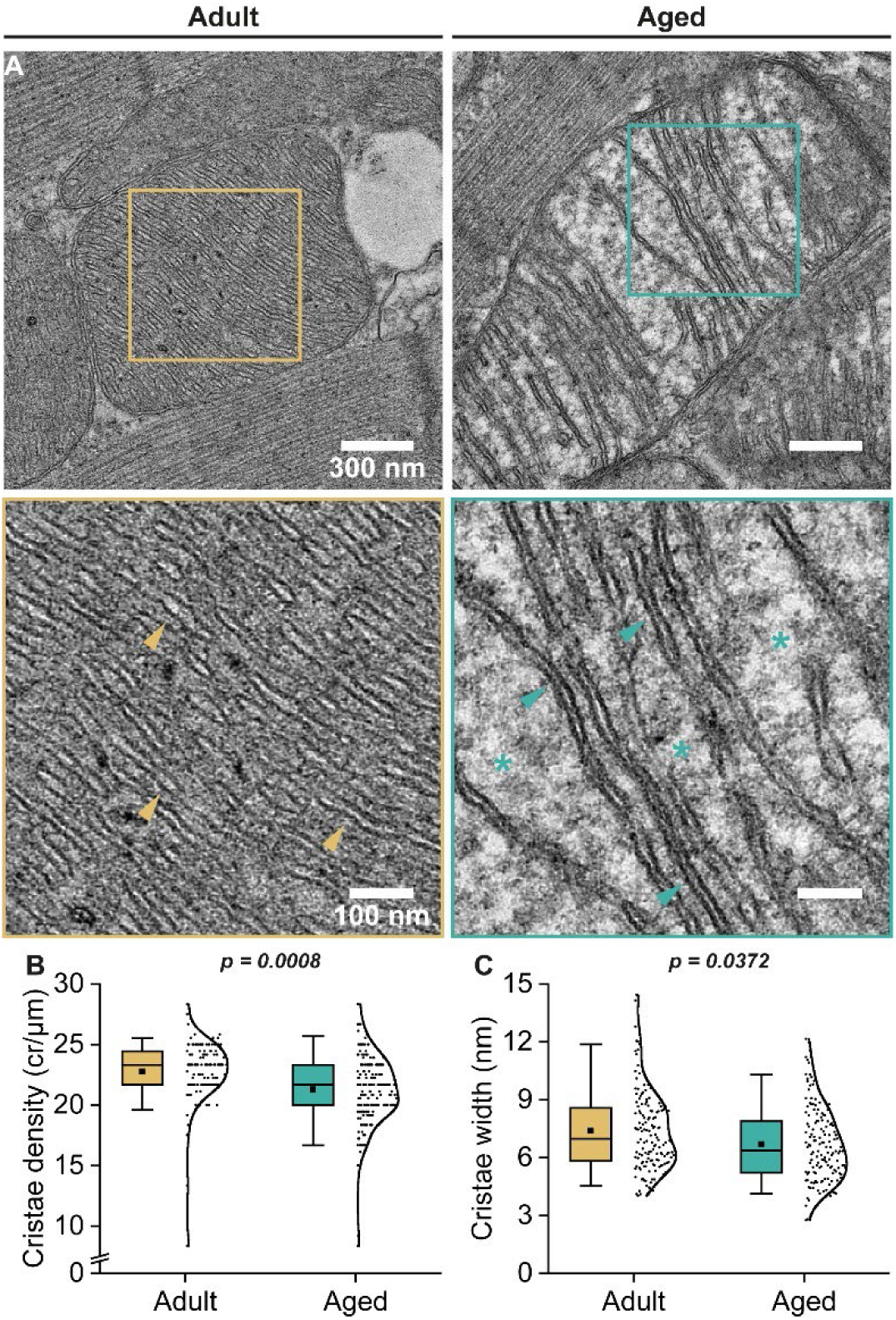
2D TEM shows a reduction cristae density and width in aged mice. **(A)** Representative images of cristae morphology from adult and aged mice hearts. Asterisks highlight regions devoid of cristae. Arrowheads highlight cristae width. **(B)** Cristae density (cristae/µm). **(C)** Cristae width (nm). Samples from 3 adult and ≥3 aged animals were used per condition. Data not following a normal distribution (P<0.05 with Shapiro-Wilk test) was normalized using rankit procedure. A hierarchical mixed random intercept model was used according to (73) to determine statistical significance. The details of the hierarchical data distribution are presented in **Supplemental Figure 1D**. For box plots: box: P25 and P75; whiskers: P5 and P95; line: median; square: mean. Every point represents a single mitochondrion.

Next, we analyzed the three-dimensional reconstruction of cardiac tissue using DeepMIB (34), a deep-learning approach that can be trained to segment the regions devoid of cristae (**Supplemental Figure 2F and Supplemental Figure 3C**). This method allowed us to segment empty or cristae-free spaces inside each mitochondrion. Our data show that a broad population of mitochondria from aged animals presented a major increase (approximately 5-fold) in cristae-free volume density (µm^3^/µm^3^) (**Figure 5**), indicating a concomitant decrease in cristae density at mitochondrial population level, as seen in human samples (**Figure 1, B and E**).

**Figure 5.**
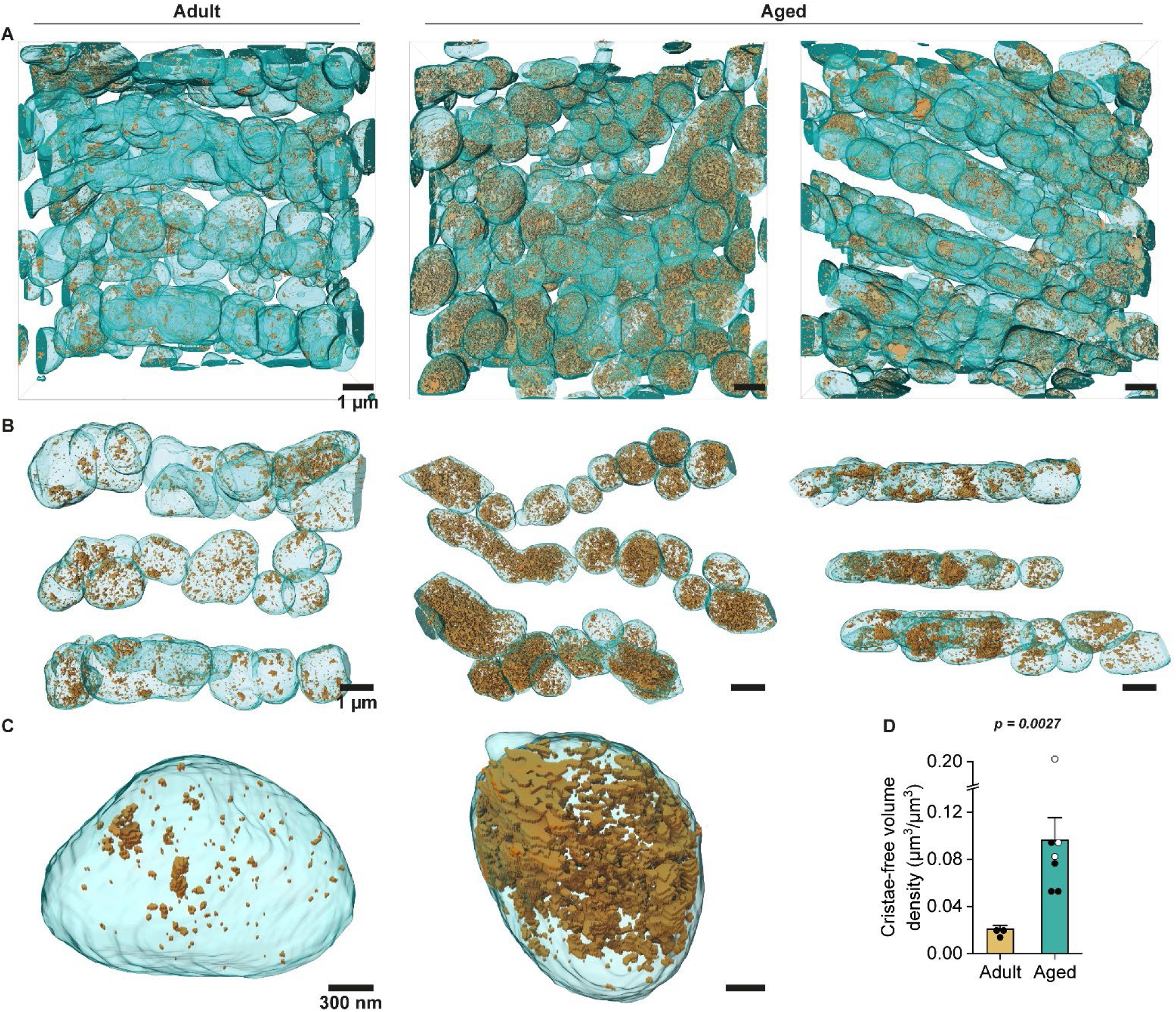
3D SBF-SEM shows that mitochondria increase cristae-free volume in aged mice hearts. **(A-C)** 3D rendering of adult and aged mice heart cristae-free volume after DeepMIB training represented as the whole sub-volume **(A)**, rows of mitochondria **(B)**, and individual mitochondria **(C)**. Mitochondria are depicted in cyan, while cristae-free volume is represented by golden structures. Samples from 1 adult (3 subsamples) and 2 aged (3 and 4 subsamples) animals were analyzed. **(D)** Cristae-free volume density (µm^3^/µm^3^), defined as the total volume of cristae-free space divided by the total mitochondrial volume in each stack. For bar plots: mean ± SEM. Every point represents a sub-volume (18 and 19 months represented as white and black points, respectively).

To further characterize three-dimensional changes in cristae ultrastructure at a higher resolution, we analyzed a mitochondrion from adult and aged mice with 3D electron tomography by using a 200 nm thin section (**Supplemental Video 3 and 4**, for adult and aged condition, respectively). To do this, we established multiple morphological parameters such as type of cristae, cristae connectivity, matrix continuity, cristae junctions, and cristae alignment (**Supplemental Figure 4**). First, we observed that cardiac IMF exhibited lamellar cristae (**Figure 6A and Supplemental Figure 5**), the classical morphology found in this type of mitochondria (20, 35, 36). These were connected by lateral nanotubules, positioned either at the middle or end of the lamella, leading to a subset of interconnected stacks (**Figure 6B**). Such interconnections are thought to allow for diffusion of solutes between cristae (35). In addition, cristae displayed membrane interruptions known as fenestrations (**Figure 6C and Supplemental Figure 5**), which allow for the diffusion of solutes within matrix sub-compartments. It is also suggested that fenestrations provide mechanical support and accommodate ATP synthase oligomers (35, 37). Lastly, cristae displayed one or multiple cristae junctions at either side (**Figure 6D**), structures that regulate the diffusion of solutes between the intermembrane space and the crista lumen (37). Regarding the age-related differences between groups, we report a decrease in the number of lamellae that comprise an interconnected stack (Figure 6B), suggesting a decrease in cristae connectivity in cardiac mitochondria from aged mice. In parallel, we found an increase in the size of cristae fenestrations (**Figure 6C and Supplemental Figure 5**), which might be part of the reason we see regions devoid of cristae in the SBF-SEM data (**Figure 3, A-C**). Changes in cristae fenestration size could imply either an increase in matrix solutes’ diffusion, a decrease in cristae surface, and, importantly for function, spatial reorganization of ATP synthase oligomers with age.

**Figure 6.**
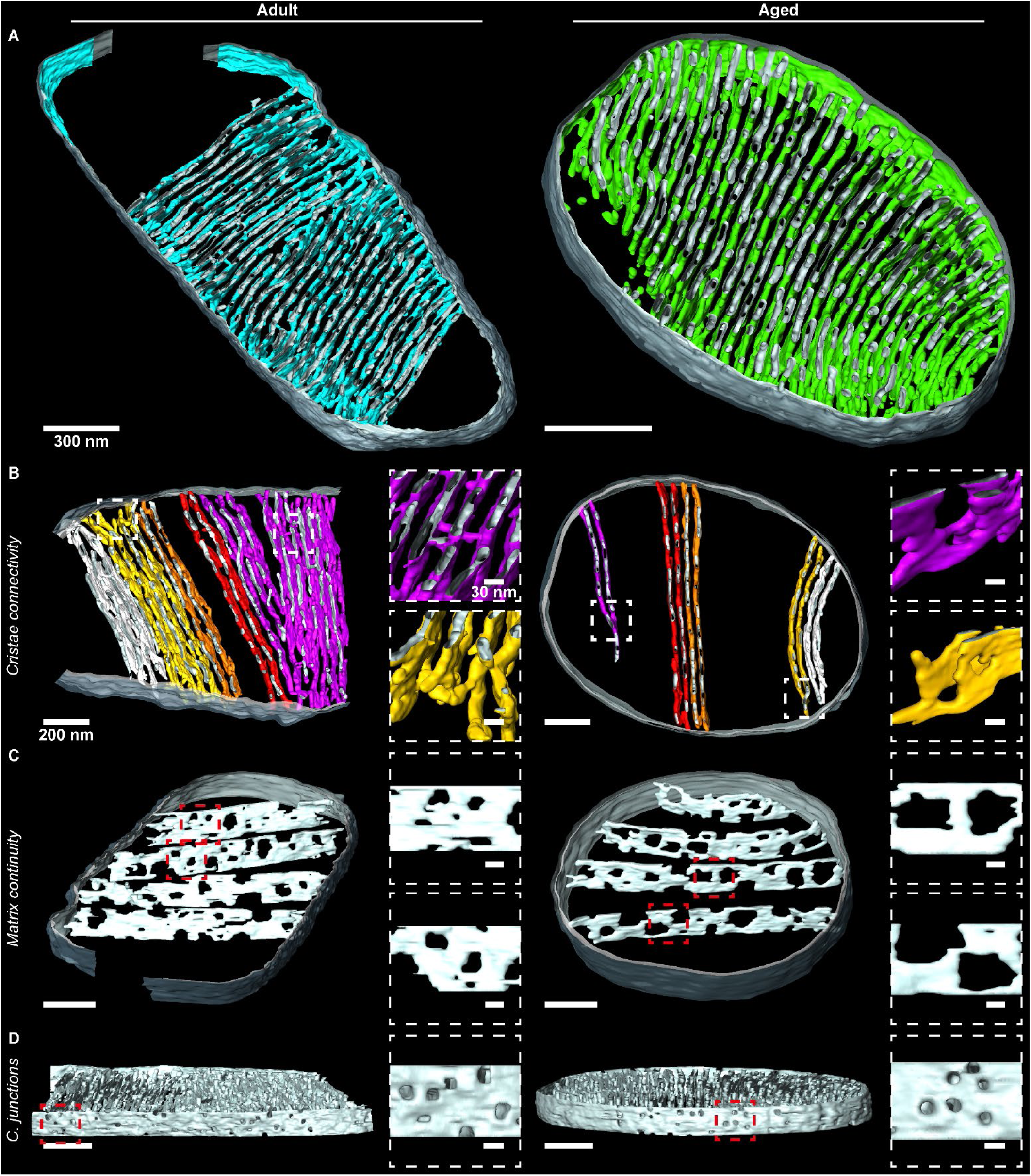
3D electron tomography shows that mitochondria decrease cristae connectivity and increase cristae fenestration size in aged mice hearts. **(A)** 3D reconstruction of cardiac mitochondria from adult and aged mice. Outer mitochondrial membrane is depicted in grey, while inner mitochondrial membrane in cyan and green. **(B-D)** Same mitochondrion as in **(A)**, highlighting cristae connectivity **(B)**, matrix continuity **(C)**, and cristae junctions **(D)**. **(B)** Cristae connectivity is depicted as substacks of connected cristae with the same color. **(C)** Matrix continuity is depicted as interruptions in lamellar cristae. **(D)** Cristae junctions are depicted as lateral openings of the cristae. Tomograms were acquired from 200 nm sections at 28000X with a voxel size of 4.10×4.10×4.10 nm^3^ (adult) and 2.05×2.05×4.10 nm^3^ (aged).

### Mitochondrial meso- and nano-dynamics machinery alterations in aged mice hearts

To evaluate if the expression of the mitochondrial meso- and nano-dynamics machinery was altered in aged mice, we determined the transcript levels of multiple genes by RT-qPCR (Figure 7A). These results showed significant differences in the expression of *Opa1* and *Drp1*, both decreased in aged mice. To confirm if these results were consistent at the protein level, we examined them by Western blot (**Figure 7, B and C**). Indeed, a reduction in Opa1 was also present at the protein level, but this was not the case for Drp1, which showed no differences. Other proteins involved in fusion, such as Mfn1 and Mfn2, were unaltered in the aged animals. Similarly, we did not find differences in fission proteins, such as Mid51, Fis1, and Mff, nor in the ratio of pro-fission (pS616) and anti-fission (pS637) Drp1 (38). Finally, we examined MICOS proteins Mic60, Mic19, and Mic10, which remained unaltered in aged mice. PGC-1α and VDAC1 levels were unaltered (**Figure 7 and Supplemental Figure 6**), suggesting no differences in mitochondrial biogenesis.

**Figure 7.**
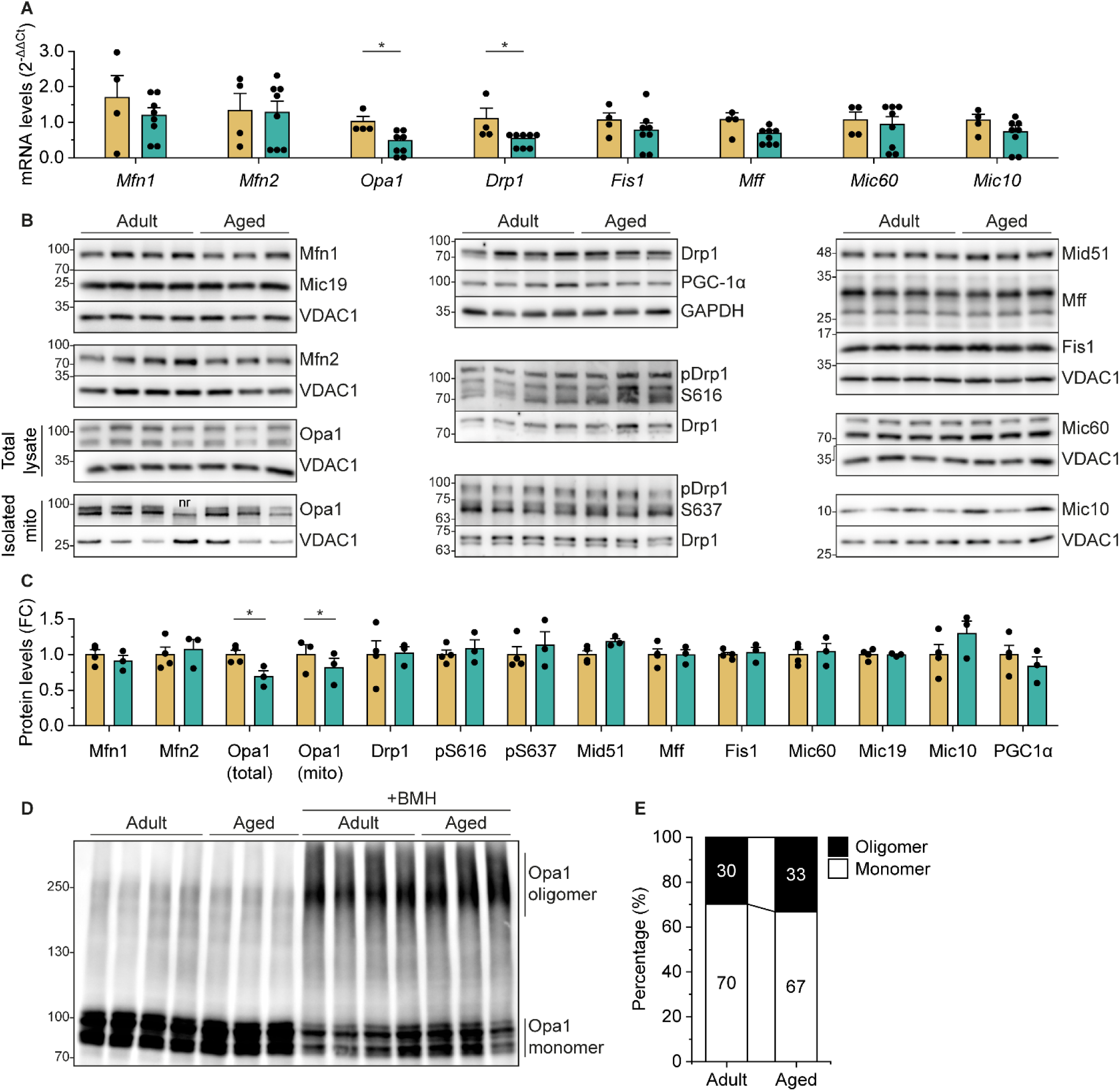
Opa1 is downregulated in aged mice hearts. **(A)** mRNA levels expressed as 2^−ΔΔCt^ against GAPDH. **(B)** Western blot of mitochondrial dynamics proteins. **(C)** Protein levels from **(B)** expressed as fold change (FC). Mitochondrial proteins were compared against VDAC1, cytosolic proteins (Drp1 and PGC-1α) were compared against GAPDH, and phosphorylated proteins (pDrp1) were compared against the corresponding total protein (Drp1). **(D)** Opa1 oligomerization assay using the chemical crosslinker BMH. **(E)** Quantification of membrane presented in **(D)** expressed as percentage. For RT-qPCR, samples from 4 adult and 8 aged animals were analyzed. For WB and Opa1 oligomerization assay, samples from 4 adults and 3 aged animals were analyzed. Data following a normal distribution (P>0.05 with Shapiro-Wilk test) was analyzed with a t-Student test to determine statistical significance. Otherwise, a Mann-Whitney test was used. (*P<0.05). For bar plots: mean ± SEM. Every point represents a mouse. nr: non-related.

The oligomerization of Opa1 controls the width of the cristae junction (9, 15) and cristae lumen (11). Since we detected a narrowing of the cristae in the aged group (**Figure 5C**), we hypothesized whether those changes were related to the degree of Opa1 oligomerization. We treated protein extracts from young and old mice with the chemical crosslinker bismaleimidohexane (BMH) (**Figure 7D**) and found that the proportions of Opa1 monomer and oligomer were comparable between the two groups. These findings suggest that cristae narrowing is independent of the oligomeric state of Opa1 (**Figure 7E**).

Our results show that age-related changes in mitochondrial and cristae ultrastructure are correlated with Opa1 downregulation, both at transcript and protein levels, but not at the oligomerization state, while other components of the fusion and fission machinery remained mostly unchanged.

### Decreased OXPHOS activity is not explained by alterations in OXPHOS composition in aged mice hearts

The changes observed in cristae ultrastructure suggested alterations in oxidative metabolism. Aging has been linked to a decrease in ATP output and an imbalance in mitochondrial bioenergetics, as well as an increase in ROS production (7). We studied mitochondrial respiration by high-resolution respirometry in freshly isolated mitochondria from adult and aged mice, in the presence of complex I substrates, malate and pyruvate (**Figure 8A**). We found no significant differences in ADP-stimulated respiration (state 3), or upon addition of ATP synthase inhibitor, oligomycin. In contrast, we found a significant reduction in the maximum oxygen flux in the noncoupled state of mitochondria from aged mice hearts compared with the adult condition (**Figure 8B**). This reduction was not explained by differences in the isolation process since mitochondrial proteins levels were comparable in both groups (**Figure 8C**).

**Figure 8.**
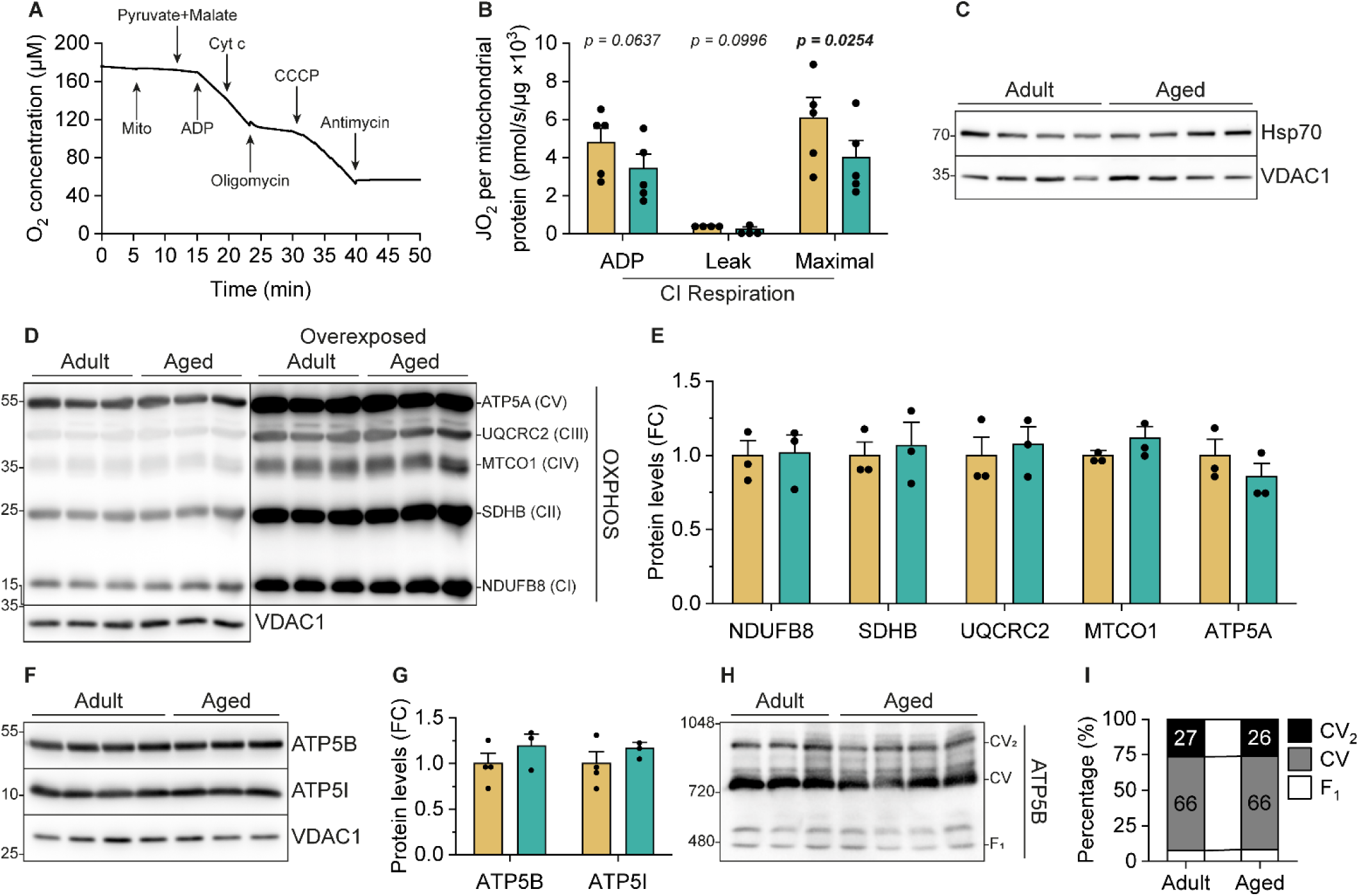
Decreased OXPHOS maximal respiration is not explained by alterations in OXPHOS subunits nor ATP synthase assembly in aged mice hearts. **(A)** Schematic representation of respirometry assay performed in Oxygraph-2k (Oroboros Instruments). Arrows represent the addition of mitochondria, substrates or inhibitors. **(B)** Oxygen consumption rate (JO_2_) of isolated mitochondria from heart left ventricle. ADP-stimulated respiration is obtained by the addition of substrates for CI (pyruvate and malate) plus ADP. Leak respiration is obtained by the addition of oligomycin. Maximal respiration is obtained by successive additions of the uncoupler CCCP. **(C)** Loading control for **(B)**. **(D)** Western blot of OXPHOS subunits. **(E)** Protein levels from **(D)** expressed as fold change (FC), normalized against VDAC1. **(F)** Western blot of ATP synthase subunits ATP5B and ATP5I. **(G)** Protein levels from **(F)** expressed as fold change (FC), normalized against VDAC1. **(H)** BN-PAGE of isolated mitochondria from adult and aged left ventricle depicting ATP synthase dimers (CV2), monomers (CV) and F1 subunit. **(I)** Quantification of membrane presented in **(H)** expressed as percentage. For respiration assays, samples from 4 animals were analyzed per condition. For WB and BN-PAGE, samples from 3 adult and 4 aged animals were analyzed. t-Student test was used to determine statistical significance.

Since the maximal, or noncoupled respiration was hindered, we searched for molecular mechanisms that could explain this reduction. First, we evaluated the level of different core subunits from the OXPHOS complexes, not finding differences between conditions (**Figure 8, D and E**), suggesting that the levels of the OXPHOS complexes remained unaltered. Next, we evaluated if the assembly of ATP synthase is altered. Given that this complex is formed by the subcomplexes Fo and F1, we measured the subunits e and β from these subcomplexes, respectively. Particularly, these proteins remained unaltered (**Figure 8, F and G**), suggesting that there are no changes in the formation of each subcomplex. Furthermore, ATP synthase can assemble in monomers, dimers or oligomers (39). To test if there were changes in the ratio of these forms, we used BN-PAGE followed by immunostaining, detecting no differences in the assembly between our adult and aged conditions (**Figure 8, H and I**).

These data indicate that the decline in noncoupled mitochondrial respiration observed in aged hearts is not attributable to changes in the expression of OXPHOS complex subunits or the levels and assembly configuration of ATP synthase. However, these findings suggest that cardiac mitochondria in aged mice might have reached a stress-induced threshold, which becomes evident under maximal conditions. Then, the loss of maximal capacity may render them more susceptible to cumulative chronic exposure to environmental insults, characteristic of aging.

## DISCUSSION

In this work, we provide an in-depth study on the effect of aging on heart mitochondrial and cristae ultrastructure in human patients and a mice model on the meso- and nanoscale. Most strikingly, we found changes in the IMM organization, unveiling alterations in cristae abundance and structure in both models.

Mitochondrial morphological abnormalities are commonly observed in aging and age-related diseases across different tissues and species, such as cardiomyopathy (2, 40). These changes are a common feature of heart failure, a condition strongly linked to aging (41), and have been observed in human samples and rat models of heart failure (41–45). In our aged human samples, we observed a significant increase in the roundness of IMF, while their area remained unchanged. This suggests that they have lost their normal rod-shaped morphology, which could be explained by the effect of sarcomere destabilization across the tissue. In agreement with this, aging is associated with changes in contractile proteins such as an increase in βMHC levels and troponin-I phosphorylation (46). In our aged mouse samples, we found that mitochondrial morphology remained generally unaltered, with a down-shift in mitochondrial volume and surface. In contrast, a study comparing 3-months and 2-year-old mice showed a decrease in mitochondrial size with an increase in mitochondrial number, leading to constant density (24), which indicates fragmentation of the mitochondrial network as opposed to our results. Additionally, they show an increase in circularity (24), while another study comparing 4-6 months and ≥20 months mice showed a decrease in mitochondrial length (23). Overall, this suggests that fragmentation is not an early event in the aging process. Likewise, studies in *Caenorhabditis elegans* showed that neither mitochondrial area, circularity nor length can be used as predictors of lifespan (47).

Previous works proposed that the mitochondrial network in striated muscle could work as an energy distribution system (48), ensuring communication across long distances, and yet being capable of segregating localized damage, protecting the overall function of the network (49). The authors proposed the inter-mitochondrial junctions as platforms for maintaining network communication (49). Moreover, close juxtaposition between cardiac mitochondria supports mitochondrial fusion, which is inhibited in alcoholic rats (50). In our aged mice, we detected that IMCS remained robustly established, in contrast with other groups that found a decrease in the level of branching of the mitochondrial network when comparing 3-months and 2-year-old mice (24).

On the nanoscale, we detected a decrease in cristae density both in human and mice cardiac mitochondria of aged individuals, suggesting a conserved mechanism in response to the chronic stressors associated with heart aging (5–7). This is consistent with other groups that have reported a reduction in cristae density in aged mice hearts (23, 24). Also, in an in vitro senescence cell model based on doxorubicin treatment, we previously showed that rat NVCM and hiPSC-derived cardiomyocytes presented an abundance of organelles with interrupted cristae and a decrease in cristae density, respectively (22). The reduction in cristae density represents a shortage of the key bioenergetic platforms that host the OXPHOS complexes needed for ATP synthesis, which is reflected in a decrease in mitochondrial maximal respiration in aged mice. We also observed a decrease in cristae width in aged mice, but not in humans. Diverse conditions lead to narrower cristae, such as starvation (11), stimulation of mitochondrial respiration by oxoglutarate (51), or glycolysis inhibition by 2-deoxy-D-glucose (13). Therefore, the narrowing of the cristae observed could represent a metabolic adaptation to enhance OXPHOS efficiency (38) given the decrease in cristae density. Indeed, a study shows that controlled overexpression of Opa1 results in narrower cristae, which conferred protection against ischemia-induced cardiac injury (52), suggesting that this remodeling might protect mitochondrial activity. Remarkably, the human aging heart displays higher dependence on glycolysis (5), which might explain why we did not detect major adaptations in cristae width in the hearts of human individuals.

Next, we found that both mitochondria analyzed by 3D electron tomography exhibited lamellar cristae. This indicates that overall morphology remains preserved, as we did not find differences in the type of cristae. However, we detected that cristae presented fewer connections between lamellae, along with an increase in the size of cristae fenestrations. These differences have not been found by others (20), likely due to the use of isolated mitochondria. Both processes could represent adaptation mechanisms to impaired OXPHOS. On one side, the decrease in cristae connectivity could isolate membrane potential inside each crista to protect neighboring cristae from depolarizing insults (53), at the cost of a slower membrane potential equilibrium across the cristae network (37). On the other hand, the apparent increase in fenestration size is likely due to an increased ATP synthase oligomer density in this nanodomain, despite the fact that the dimerization levels of ATP synthase were not changed. This is supported by data from our group that shows larger cristae fenestrations in senescent rat NVCM mitochondria, alongside a decrease in ATP synthase mobility in senescent hiPSC-derived cardiomyocytes, which is associated with ATP synthase oligomerization but a decrease in ATP production (54). Overall, these changes could be explained by cristae remodeling events, although their kinetics remain unknown and its correlation with ATP synthase organization demand further investigation.

Abnormal mitochondrial morphology may result from impairments in mitochondrial meso-dynamics. Cardiac defects and premature death can be observed in a murine Drp1 knockout, and Mfn1/Mfn2 double knockout (55), while mice Mfn1/Mfn2/Drp1 cardiac triple knockout showed a longer survival but accumulated abnormal mitochondria (56), similar to the results obtained in an Mff/Mfn1 double knockout model (57). We analyzed the expression of the mitochondrial meso-dynamics machinery in aged human right atria and aged mice left ventricle, finding mostly no differences in transcript nor protein levels, in alignment with our findings in doxorubicin-induced senescence cell models in rat NVCM and hiPSC-derived cardiomyocytes (22). This is inconsistent with studies in age-related cardiovascular diseases, such as heart failure and cardiomyopathies, where mitochondrial meso-dynamic adaptations seem to be case-dependent. For instance, in the human failing heart, MFN2 protein levels decreased, whereas DRP1 and FIS1 protein levels augmented (58). Another study reported elevated levels of MFN1, MFN2, and DRP1 in explants of patients with ischemic or dilated cardiomyopathy (42). Additionally, impaired mitochondrial fusion in alcohol-fed rats was linked to Mfn1 downregulation (50). However, as we worked with a model in early stages of aging, and without serious cardiac impairments, this inconsistency could point to the hypothesis that alterations in protein expression occur in advanced stages of aging. Although we found that the meso-dynamics machinery remained mostly unaltered, we detected downregulation of mitochondrial fusion protein Opa1 in our aged mice. This is consistent with a decrease in Opa1 levels in aged cardiac samples from Tibetan sheep (21) and in skeletal muscle samples from aged sedentary individuals(59). Meanwhile, in a skeletal muscle-specific Opa1^−^/^−^ inducible mice model, the animals developed precocious aging (59). In addition, in heart failure, human and rat cardiac samples presented decreased Opa1 protein levels, while mRNA levels remained unchanged, suggesting a post-translational regulation of Opa1 upon cardiac stress-induced adaptation (42). It has been shown that defects in Opa1 can inhibit mitochondrial fusion, changing the balance of mitochondrial meso-dynamics towards a fragmented network (60). In early heart aging, this seems not to be the case, given that we do not see any changes in mitochondrial size, however, in late heart aging other authors have reported a fragmented network (24). This suggests that Opa1 downregulation does not have a short-term effect, but a long-term effect in meso-dynamics. Altogether, our data contributes to a body of literature highlighting the relevance of Opa1 in cardiac mitochondrial and heart homeostasis. Moreover, these studies suggest that in cardiac dysfunction and in the aged heart, mitochondrial meso-dynamics controlling proteins distinctly adapt, but morphology adaptations are not always accompanied by protein level modifications, although post-translational modifications that control the dynamics mechanisms cannot be excluded(38).

In parallel to meso-dynamics, Opa1, MICOS, and ATP synthase are essential components of the mitochondrial nano-dynamics machinery. Particularly, Opa1 determines cristae abundance and shape (9, 16, 27). As previously said, we found Opa1 downregulation in the aged mice hearts, but no changes in proteins from the MICOS complex, or the expression and assembly of dimeric forms of ATP synthase. This is consistent with studies in heterozygous Opa1^+^/^−^ mice, where at 6 months the heart showed cristae abnormalities (61), and at 12 months there was cristae loss (62). Altogether, this indicates that Opa1 downregulation is responsible for the decrease in cristae density found in aged mice. In contrast, this would not be due to lower levels of the MICOS complex, as others suggest (24). In addition to cristae maintenance, Opa1 regulates cristae width depending on its oligomerization state, which can be controlled by cell (63) and metabolic cues (11) to modulate OXPHOS activity by altering substrate concentration (64) and the assembly of supercomplexes (9, 10, 65). According to our results in cristae narrowing in the aged mice cardiac mitochondria, we expected an increase in the oligomeric form of Opa1, however, we did not find differences between both groups in mice. Thus, we suggest that other mechanisms may be responsible for cristae narrowing in the mitochondria of aged mice, such as a decreased ETC activity and membrane energization (22).

While the impact of Opa1-linked reduction in cristae density was not observable in ADP-stimulated and leak respiration, which remained unchanged between groups, we found a drop in maximal respiration. This is consistent with studies in heterozygous Opa1^+^/^−^ mice, where at 12 months there was reduced maximal respiration, leading to the development of cardiomyopathies (62). In contrast, others did not detect differences in this state when comparing total heart-isolated mitochondria from 20-week versus 80-week old mice (20). However, in our study, we isolated the left ventricle, therefore suggesting that differences in mitochondrial respiration may depend on the specific section of the heart selected for analysis. Additionally, differences could be explained by the effect of particular mitochondrial subpopulations. Our data highlights crista alterations in IMF mitochondria, which is consistent with previous reports, where this population showed impaired OXPHOS capacity (5, 23, 31).

Interestingly, the differences in maximal respiration were not explained by differences in the protein levels from the OXPHOS complexes. This was surprising because, if cristae abundance decreases, one would expect that the proteins associated would also decrease. Instead, the levels of the OXPHOS complexes remained constant, and as they are located in this subdomain, it is implied that every crista accommodates more OXPHOS complexes than before. In addition, ATP synthase dimerization neither explains the difference in maximal respiration. This contradicts results from others, in which they detected a decrease in the abundance of dimeric and oligomeric forms of ATP synthase in old mouse hearts (23). Interestingly, the authors did not find differences in the levels of ATP synthase subunits, similar to our results. However, it needs to be taken into consideration that both works differ in the source from which mitochondria were isolated, and the mitochondrial subpopulation studied. Alternatively, Alternatively, it could also be related a drop in supercomplexes assembly, caused by the partial loss of OPA1 (66), as seen by others in the rat’s aging heart (67).

Physiologically, a decrease in maximal respiration could be interpreted as a decline in the capacity to respond to stressors, which leads to a decrease in the ability to answer higher metabolic demands as in exercising (68), which goes in line with defects seen in the aging heart (6). We speculate that the proposed increase in OXPHOS complexes density in the cristae optimizes the conditions for normal respiration but cuts the possibility of enhancing OXPHOS under stress. This is equivalent to a loss of OXPHOS system plasticity of mitochondria in the aging heart. As seen in mice hearts, this is accompanied by cardiac hypertrophy, suggesting that as a tissue, the heart responds remodeling to fulfill its demands as mitochondrial capacity is insufficient. Interestingly, cristae alterations were previously reported in cardiac hypertrophy (69, 70). Finally, lower mitochondrial maximal respiration in aged hearts might also represent a minor stress-response capacity, that would promote the mitochondrial permeability transition pore and subsequent cardiac cell death as widely reported in cardiac ischemia-reperfusion and acute infarct (71).

In conclusion, our study reveals critical insights into the mitochondrial alterations that accompany cardiac aging, particularly changes in cristae abundance and structure. While mitochondrial morphology abnormalities are commonly observed in aging and heart failure, our findings suggest that these changes do not necessarily involve early and massive fragmentation of the mitochondrial network. Instead, we observed the downregulation of Opa1 linked to a decrease in cristae density, and a reduction in maximal respiration in aged mice hearts. Together, these changes result in a decline of OXPHOS system plasticity, which could precede more overt signs of mitochondrial dysfunction, making it a potential early indicator of cardiac aging and the onset of heart failure, even before structural changes in mitochondrial size or network fragmentation become evident. As cristae are crucial for maximizing the surface area available for OXPHOS, a reduction in cristae width may temporarily increase the efficiency of energy metabolism, but under high-demand conditions such as exercise, it becomes compromised due to a lack of further capacity. Consequently, compounding stresses of aging, such as comorbidities, contribute to the age-related decline in cardiac function.

## MATERIALS AND METHODS

### Sex as a biological variable

Sex was not taken into account as a biological variable in this study, which was conducted exclusively on male mice and humans. Therefore, the relevance of these findings in female remains unknown.

### Human samples

Heart tissue from male patients over 49 years old referred to elective myocardial revascularization surgery was collected following the Guide for Approval of Research Involving Human Subjects or Use of Human Specimens (#180913005), approved by the Ethics Review Commitee from Pontificia Universidad Católica de Chile Faculty of Medicine. A total of 12 participants were included for this study, not considering the patients that: (a) Presented an emergency revascularization or pre-operative cardiac shock, (b) Present a history of acute myocardial infarction during 30 days prior to surgery, (c) History of neoplasms, rheumatologic disease or any active chronic inflammatory condition, (d) Chronic steroid treatment, (e) Untreated thyroid disease or (f) Intercurrent infections. Samples were collected after obtaining an informed consent, approved and signed by the patient, before the medical team performed the operation. Human atrial samples of approximately 1 cm and 2 mm thick were separated for sample fixation for TEM, qPCR, and WB.

### Animals

Male C57BL6/J mice were bred in the bioterium of the Instituto de Nutrición y Tecnología de los Alimentos (INTA) at Universidad de Chile. All studies were conducted following the Guide for the Care and Use of Laboratory Animals (#220818001) and were approved by the Ethics Review Committee. The animals were euthanized by cervical dislocation and their hearts were collected. Cardiovascular surgeons from the Hospital Clínico de la Universidad Católica carefully dissected the heart into the four cavities plus the septum. The left ventricle was subdivided into two parts – the apex and the ventricular wall– which were used for biochemical or electron microscopy assays, respectively.

### Transmission electron microscopy (TEM)

For TEM, whole tissue segments (left ventricle apex from mice or right atrial appendage from human samples) were fixed in 2.5% glutaraldehyde (Science Services, #E16216) in 0.1 M cacodylate buffer pH 7.4 (Sciences Services #11652) at room temperature. Subsequently, they were washed in 0.1 M cacodylate buffer pH 7.4 and postfixed in 1% osmium tetroxide (Science Services, #19150) in 0.1 M cacodylate buffer pH 7.4 at room temperature for 2 hours. Finally, the samples were dehydrated stepwise in a graded acetone series and embedded in Epoxy Embedding Medium (Sigma Aldrich #45359). Sections of 90 nm for 2D classical analysis or 200 nm for tomographs were obtained using the ultramicrotome EM UC7, Leica Ultracut R, and then stained for 4% uranyl acetate in methanol for 1 minute and Reynold’s lead citrate for 4 minutes (Leica #16707235). Images of the sections and tomograms were acquired in the transmission electron microscope TALOS F200C G2 system (Thermo Scientific) equipped with a Ceta 16M CMOS camera, at 200 kV, at the Unidad de Microscopía Avanzada de la Pontificia Universidad Católica (UMA-UC).

### Mitochondrial morphology analysis

For 2D morphometric analysis, 8500X and 28000X images were obtained and analyzed using ImageJ. Images at 8500X were used to measure area and roundness through manual segmentation. Images at 28000X were used for analysis of cristae morphology. To measure cristae density, we established a simple method where a series of 200 nm lines was traced at different sites of the mitochondrion. We then counted the number of distinguishable cristae crossing these lines and calculated an average per mitochondrion. To measure cristae width, 1 to 4 cristae per mitochondrion were selected to calculate the average of 3 measurements per cristae.

### Electron tomography

Single-tilt image series (±60°, 1° increment) were acquired using a 512×512 or 1024×1024 detector array. Tomograms were aligned using Inspect 3D software, applying three algorithms: Weighted Back Projection, Simultaneous Iterative Reconstruction Technique, or Expectation Maximization. Aligned tomograms were manually segmented using Amira.

### SBF-SEM sample preparation

After primary fixation in glutaraldehyde (2.5% in 0.1M cacodylate buffer, pH 7.4; Science services, #E11652), left ventricular apex biosections were shipped to the Integrated Bioimaging Facility (IBiOs) in Osnabrück, Germany, were the preparation and imaging of the sections was performed. To ensure pronounced contrast and stability necessary for blockface imaging, the tissue was prepared via an adapted version of the Ellisman protocol(72).

In detail, after washing the samples in buffer cacodylate 5 times for 7 minutes, they were post-fixed with both 2% osmium-tetroxide (Science Services, #19150) and 1.5% potassium ferrocyanide (Science Services, #E11652) in 0.1 M cacodylate buffer simultaneously for 80 minutes on ice with gentle agitation. Following that, all steps were performed in-between rounds of washing with double-distilled water, repeated 5 times for 7 minutes with gentle agitation, at room temperature, unless stated otherwise. Following initial osmication, samples were treated with aqueous 1% thiocarbohydrazide (Sigma-Alrich, #223220-5G) for 20 minutes. Consecutively, secondary osmication was performed with 2% osmium in water for an additional 30 minutes, after which samples were incubated in 1% uranyl acetate-dihydrate (EMS, #22400) in water over night at 4°C.

The next day, samples were treated with Walton’s lead aspartate for 40 minutes at 60°C aspartate (Lead(II) nitrate (Carl-Roth, #10099-74-8), L-Aspartate (Serva, #14180.02), KOH (Merck, #109112)), washed with pre-warmed water, before undergoing a graded dehydration series in aqueous ethanol mixtures. Each step (20%, 50%, 70%, 90%, 2× 100%) was performed for 7 minutes on ice, after which replacing the ethanol with ice-cold acetone for 10 minutes at room temperature. Consecutively, the acetone was replaced by room-temperature, water-free acetone (VWR Chemicals, # 83683.23) for additional 10 minutes, before infiltrating in fresh Epon 812 (Sigma Aldrich, #45359-1EA-F; hard formula) with increasing concentrations of 25%, 50% and 75%, for 2 hours each, followed by an overnight incubation in 100% Epon. On the next day, tissue samples were transferred into fresh resin, containing 1.5% Ketjenblack (w/w) (TAAB, #C409) for additional 4 hours, until polymerized at 60°C for 48 hours.

Once cured, sample blocks were trimmed to approximately 300 µm³ blocks and glued onto aluminum rivets with two-component silver epoxy adhesive (Microtonano, #AG29D). Sample stubs were then trimmed further into a frustum shape, for preparation of toluidine blue sections, which were helpful to identify orientation of the tissue block, assessing sample integrity, as well as pre-selecting areas suitable for ultrastructural analysis. Samples were then additionally grounded by coating with a 20 nm thin gold layer, before being inserted into the stage of a 3View2XP (Gatan, Pleasanton, CA, USA), fitted inside of a JSM 7200F (JEOL Ltd., Tokyo, Japan).

Large-scan overviews helped in finding suitable regions for further high-resolution image series acquisition. Regions of interest were chosen by identifying cardiomyocytes approached in a longitudinal cross-section, with clearly observable Z-lines, I-bands, and evenly-spaced sarcomeres. This focused analysis on regions with high densities of IMF—while excluding nuclei, intercalated discs, and sarcolemma—ensured that subsequent measurements accurately reflected mitochondrial ultrastructure.

### Image acquisition

The tissue samples proved to be stable under imaging conditions with 3.1 kV accelerating voltage, 30 nm condenser aperture, high vacuum mode of 10 Pa, under the addition of a positive stage bias of 600 V. Final parameters for high resolution imaging were set to voxel size of 2 x 2 x 25 nm, with a dwell time of 2.1 µs. Overall, a volume of approximately 20 x 20 x 6 µm were acquired, corresponding to 250 consecutive images, sized 10240 × 10240 pixel. The image acquisition and stage were controlled via Gatan Digital Micrograph (Version 3.32.2403.0).

### SBF-SEM segmentation and data acquisition

Individual slices were processed in Microscopy Image Browser (MIB) (33) segmenting and visualization software. The aligned, denoised, 8-bit converted and cropped image volumes were then exported and binned whenever possible, to reduce computational resources to a minimum.

For extracting the mitochondrial context, SBF-SEM volumes were segmented using Empanada (32) following the instructions provided by the developers. Empanada is a napari plugin that uses MitoNet for panoptic segmentation of mitochondria, also offering a tool to train and finetune models based on sample characteristics. Each dataset was first anonymized, then segmented by a dedicated model previously trained utilizing the Empanada finetune module. The Segmentation Confidence threshold was 0.5 or lower, while the center confidence threshold was set to 0.05, except for two datasets from the 18-month samples, in which it was set to 0.3. In our experience lowering these values significantly improved over segmentation by increasing the number of objects detected. The resulting segmentation model was saved as a TIF file and visualized using MIB.

For consecutive analysis and statistical read-out, the correct splitting of mitochondria in three dimensions was important. Since fusion and fission, particularly in clusters of IMF, made automatic panoptic segmentation error-prone, a semi-automatic curation approach using the 3D object splitting tool in MIB was performed; in MIB: Tools/Object Separation/3D, volume). Additionally, all datasets were manually revised. The final model files were binned to isometric voxel size and then analyzed for volume and surface area.

For the analysis of IMCS, final panoptic labels were transformed into a semantic model, which was then used for main and secondary organelle targets in the MIB MCcalc plug-in for calculation of contacts and distance distributions; in MIB: Plugins/Organelle/Analysis/MCcalc). Overall, the probing distance of extended rays were 200 nm, with a contact cut off after 100 nm, effectively selecting contact points between each mitochondrion within the range of 100 nm in 2D. The result is the detection of two juxta positions in which mitochondria are next to each other, which does not properly reflect the definition of membrane contact sites, as a planar sheet between two objects. To address this, the detected contacts were expanded by 50 nm to fuse the facing membranes, and then the original mitochondria mask was subtracted from the now formed inter-mitochondrial space. Additionally, to take into account curvature of mitochondria, the selection was smoothened in 3D (in MIB: Selection/Smooth /Selection). Finally, in order to read-out not a volume, but contact areas, the smoothened output was processed in the Surface Area 3D plug-in (in MIB: Plugins/Organelle Analysis/Surface Area3D), which calculates surface area, defined from the center lines of each cross section.

### SBF-SEM cristae quantitative data

For analysis of mitochondrial cristae organization, the mitochondria labels were used as mask in order to apply a selective threshold. By using the Otsu method, mitochondria were divided based on their mean intensity, effectively labeling electron lucent inner mitochondrial membrane space and more electron dense mitochondrial membrane material. This approach relies heavily on intensity distribution of pixel values alone, which can be problematic when there are significant variations in contrast or illumination across different samples and phenotypes. To reduce the variability in contrast between different conditions, prior to analysis, all datasets were normalized globally, based on the mitochondria volumes within the stacks. Still, to take into account the different phenotypes of senescent mice, simply using a threshold based on the mean intensity would falsify the result as the threshold would no longer properly reflect membrane architecture. Instead, a sub-volume of 5 random slices per dataset was segmented and then used as ground truth for training a deep learning model Z2C + DLv3 Resnet50, with a patch size of 256×256×5 patches in DeepMIB (34). The prediction was performed only within the mitochondrial-masked area. This approach allowed for detection of spatial context of the inner mitochondrial membrane spots and reduce reliance on pixel intensity values alone, making it less susceptible to thresholding biases associated with variations in staining. The now segmented volume of IMM space was then measured and plotted.

### Western blotting

Left ventricular wall tissue was homogenized using a Bioprep 6 homogenizer with 3.0 mm zirconium pearls in buffer RIPA (Thermo Scientific, #89901), with protease and phosphatase inhibitors (Sigma Aldrich: PMSF, #P7626; leupeptin, #103476-89-7; pepstatin A, #26305-03-3 and aprotinin, #A3428, NAF, #S6776). The samples were centrifuged at 12000 g at 4°C for 10 minutes and the supernatant was collected. Protein concentration was evaluated using the Micro BCA protein assay kit (Thermo Scientific, #23235). Samples for electrophoresis were prepared with 30 or 20 µg of protein when total lysate or isolated mitochondria were used, respectively, and 4X SDS-PAGE sample buffer. Then, they were heated at 95°C, or 45°C when the OXPHOS antibodies were used, for 5 minutes. Electrophoresis gels were prepared at 8%, 10%, or 12% (w/v) polyacrylamide gels, and then transferred to PVDF membranes. These were blocked in 5% (w/v) milk in TBS 0.1% (v/v) Tween 20 solution at room temperature for 1 hour in agitation. Primary antibodies were prepared in blocking solution and incubated at 4°C overnight. After washing (3 times for 5 min each), the membranes were incubated for 1 hour with a HRP linked secondary antibody in blocking solution (1:5000). Blots were washed again, and bands were detected using SuperSignal West Dura (Thermo Scientific, #34075) in a Firereader 10V-Plus 20M Uvitec. Antibody details are available in **Supplemental Table 1**.

### Reverse transcription-quantitative PCR (RT-qPCR)

RNA was obtained using TRIzol reagent (Thermo Scientific #15596026) and cDNA was synthesized using iScript reagent (Bio-Rad #1708890) according to the manufacturer’s instructions. qPCR was performed in a QuantStudio3 (Applied Biosystems) using Kapa SYBR Fast Master Mix (Sigma Aldrich #KK4602) for mice samples and PowerUp SYBR Green Master Mix (Thermo Fischer, #A265741) for human samples, in 10 µl reaction in Fast mode. Primer details are available in **Supplemental Table 2**.

### Mitochondrial isolation

All procedures were conducted at 4°C. Left ventricle tissue was maintained in ice-cold BIOPS buffer (2.77 mM CaK_2_EGTA, 7.23 mM K_2_EGTA, 5.77 mM Na_2_ATP, 6.56 mM MgCl_2_ ·6H_2_O, 20 mM Taurine, 15 mM Na_2_Phosphocreatine, 20 mM Imidazole, 0.5 mM Dithiothreitol, 50 mM MES hydrate, pH 7.1) until extraction. Left ventricle mitochondria were isolated as previously described with some modifications (Oroboros protocol). Briefly, heart tissue was washed in ice-cold CP1 buffer (100 mM KCl, 50 mM MOPS, 5 mM MgSO4·7H_2_O, 1 mM EGTA, 1 mM Na_2_ATP, pH 7.4). The tissue was minced using a single edge razor in buffer CP1 and digested with trypsin 0.05% for 2.5 minutes. Immediately after digestion, 2 mL of buffer CP2 (CP1 plus 0.2% BSA) were added to the solution. The digested tissue was transferred to a Potter-Elvehjem homogenizer (800 rpm) and homogenized four times.

The homogenate was transferred to plastic tubes and centrifugated at 800 g at 4°C for 10 min. The supernatant was collected and centrifugated at 3000 g at 4°C for 10 min. The supernatant was discarded, and the pellet was resuspended with buffer CP2 and centrifugated again at 3000g at 4°C for 10 min. The pellet was resuspended in KME buffer (100 mM KCl, 50 mM MOPS, 0.5 mM EGTA) supplemented with protease inhibitors (Sigma Aldrich: leupeptin, #103476-89-7; pepstatin A, #26305-03-3, and aprotinin, #A3428), until mitochondrial respiration measurements. Protein concentration was quantified according to manufacturer’s instructions (Pierce™ Rapid Gold BCA, #A53225)

### Oxygen consumption

For O2k respirometry, freshly isolated mitochondria were used. 20 µg of cardiac mitochondria were added to 0.5 mL O2k chamber in Mir05 buffer (0.5 mM EGTA, 3 MgCl_2_, 60 mM Lactobionic acid, 20 mM Taurine, 10 mM KH_2_PO_4_, 20 mM HEPES, 110 mM D-Sucrose). The SUIT protocol 0006 O2 mt D047, which evaluates the N-pathway in isolated mitochondria was used, in which consequent additions of substrates were injected into the closed chamber. Final concentrations used were: 5 mM pyruvate, 2 mM malate, 2.5 mM ADP, 1.5 mM MgCl_2_, 10 µM cytochrome c, 12.5 nM Oligomycin, 3-5 µM FCCP, 2.5 µM Antimycin A.

### Blue native-polyacrylamide gel electrophoresis (BN-PAGE)

BN-PAGE was performed as previously described (PMID: 17406264). Mitochondria were isolated as previously described and quantified by Bradford Assay (ROTI Nanoquant, Roth, #K880.3). Aliquots of 50 µg of mitochondria were centrifuged at 4°C, 9000g for 10 min and the pellet was resuspended in blue native buffer with 4% digitonin (Sigma Aldrich, #D5628). A 3 – 12% polyacrylamide gradient gel (Serva, #43250) was used alongside a molecular weight marker (NativeMark, Thermo Fisher Scientific, #LC0725). The samples were transferred to a PVDF membrane for immunoblotting for Complex V visualization, using ATP5B. Bands were detected by chemiluminescence (Super Signal West PICO PLUS, Thermo Fisher Scientific, #34580)

### Statistical analysis

For TEM and SBF-SEM data, a hierarchical analysis was performed, according to Sikkel, et al. (73). Briefly, this pipeline quantified the amount of clustering between datapoints preceding from different animals and applies a correction to the statistical test. Not normally distributed data had to be converted using a rankit approach in GraphPad 9.0. P-values < 0.05 were considered statistically significant. For RT-qPCR and WB data, a non-paired t-Student or Mann-Whitney test was used depending on the distribution of the data, evaluated by a Shapiro-Wilk test. For oxygraphy data, a paired t-Student test was used. P-values < 0.05 were considered statistically significant. Graphs were plotted using OriginLab Pro 2024.

## AUTHOR CONTRIBUTIONS

VE and KBB designed the experiments. IM-R., GB., and VE wrote the main manuscript text. IM-R, GB, WG, SM conducted experiments. LB obtained SBF-SEM data. LB, IM-R., SM established the pipeline for SBF-SEM data analysis. GB established the pipeline for tomograph data acquisition and segmentation. Data analysis was done by IM-R, GB, LB, WG, SM, HV, LG-O, KBB. HV and LG-O, harvested human samples and, KBB, HV, LG-O dissected mice heart. FD-C maintained the mice colony. A. C. provided mice samples.

## ACKNOWLEDGEMENTS

We want to thank Alejandro Munizaga and Liseth Garibaldi for TEM sample preparation. We also want to thank Rodrigo Troncoso for his assistance. The study was supported by a grant from the BMBF/DLR (FKZ 01DN19046) for S.M./K.B.B., and PCI/ANID-BMBF (180060) for V.E./H.V./K.B.B,. and FONDECYT (1191770 and 1231557) for V.E and FONDECYT (1211270) for H.V. I.M.-R. was supported by ANID Ph.D. fellowships 21201041.

## REFERENCES

1. Ageing and health [Internet]. https://www.who.int/news-room/fact-sheets/detail/ageing-and-health. Accessed September 5, 2024.

2. López-Otín C, et al. Hallmarks of aging: An expanding universe. Cell. 2023;186(2):243–278.

3. Murphy E, et al. Mitochondrial Function, Biology, and Role in Disease: A Scientific Statement From the American Heart Association. Circ Res. 2016;118(12):1960–1991.

4. Martin SS, et al. 2024 Heart Disease and Stroke Statistics: A Report of US and Global Data From the American Heart Association. Circulation. 2024;149(8). 10.1161/CIR.0000000000001209.

5. Lesnefsky EJ, Chen Q, Hoppel CL. Mitochondrial Metabolism in Aging Heart. Circulation Research. 2016;118(10):1593–1611.

6. Brown DA, et al. Mitochondrial function as a therapeutic target in heart failure. Nat Rev Cardiol. 2017;14(4):238–250.

7. Yaniv Y, Juhaszova M, Sollott SJ. Age-related changes of myocardial ATP supply and demand mechanisms. Trends Endocrinol Metab. 2013;24(10):495–505.

8. Abdellatif M, et al. Hallmarks of cardiovascular ageing. Nat Rev Cardiol. 2023;20(11):754–777.

9. Cogliati S, Enriquez JA, Scorrano L. Mitochondrial Cristae: Where Beauty Meets Functionality. Trends in Biochemical Sciences. 2016;41(3):261–273.

10. Cogliati S, et al. Mitochondrial Cristae Shape Determines Respiratory Chain Supercomplexes Assembly and Respiratory Efficiency. Cell. 2013;155(1):160–171.

11. Patten DA, et al. OPA1-dependent cristae modulation is essential for cellular adaptation to metabolic demand. EMBO J. 2014;33(22):2676–2691.

12. Nielsen J, et al. Plasticity in mitochondrial cristae density allows metabolic capacity modulation in human skeletal muscle. The Journal of Physiology. 2017;595(9):2839–2847.

13. Salewskij K, et al. The spatio-temporal organization of mitochondrial F1FO ATP synthase in cristae depends on its activity mode. Biochimica et Biophysica Acta (BBA) - Bioenergetics. 2020;1861(1):148091.

14. Hackenbrock CR. Ultrastructural bases for metabolically linked mechanical activity in mitochondria.

15. Scorrano L, et al. A Distinct Pathway Remodels Mitochondrial Cristae and Mobilizes Cytochrome c during Apoptosis. Developmental Cell. 2002;2(1):55–67.

16. Kondadi AK, Anand R, Reichert AS. Cristae Membrane Dynamics – A Paradigm Change. Trends in Cell Biology. 2020;30(12):923–936.

17. Chen H, et al. Mitochondrial Fusion Is Required for mtDNA Stability in Skeletal Muscle and Tolerance of mtDNA Mutations. Cell. 2010;141(2):280–289.

18. Ban-Ishihara R, et al. Dynamics of nucleoid structure regulated by mitochondrial fission contributes to cristae reformation and release of cytochrome *c*. Proc Natl Acad Sci USA. 2013;110(29):11863–11868.

19. Daum B, et al. Age-dependent dissociation of ATP synthase dimers and loss of inner-membrane cristae in mitochondria. Proceedings of the National Academy of Sciences. 2013;110(38):15301–15306.

20. Brandt T, et al. Changes of mitochondrial ultrastructure and function during ageing in mice and Drosophila. eLife. 2017;6:e24662.

21. Wang G, et al. Effects of Aging on Expression of Mic60 and OPA1 and Mitochondrial Morphology in Myocardium of Tibetan Sheep. Animals (Basel). 2020;10(11):2160.

22. Morris S, et al. Inner mitochondrial membrane structure and fusion dynamics are altered in senescent human iPSC-derived and primary rat cardiomyocytes. Biochimica et Biophysica Acta (BBA) - Bioenergetics. 2023;1864(2):148949.

23. Bou-Teen D, et al. Defective dimerization of FoF1-ATP synthase secondary to glycation favors mitochondrial energy deficiency in cardiomyocytes during aging. Aging Cell. 2022;21(3):e13564.

24. Vue Z, et al. Three-dimensional mitochondria reconstructions of murine cardiac muscle changes in size across aging. American Journal of Physiology-Heart and Circulatory Physiology. 2023;325(5):H965–H982.

25. Vincent AE, et al. The Spectrum of Mitochondrial Ultrastructural Defects in Mitochondrial Myopathy. Sci Rep. 2016;6(1):30610.

26. Wai T. Is mitochondrial morphology important for cellular physiology? Trends in Endocrinology & Metabolism. 2024;35(10):854–871.

27. Eisner V, Picard M, Hajnóczky G. Mitochondrial dynamics in adaptive and maladaptive cellular stress responses. Nat Cell Biol. 2018;20(7):755–765.

28. Yin FC, et al. Use of tibial length to quantify cardiac hypertrophy: application in the aging rat. Am J Physiol. 1982;243(6):H941–947.

29. Bleck CKE, et al. Subcellular connectomic analyses of energy networks in striated muscle. Nat Commun. 2018;9(1):5111.

30. Courson JA, et al. Serial Block-Face Scanning Electron Microscopy (SBF-SEM) of Biological Tissue Samples. J Vis Exp. [published online ahead of print: March 26, 2021];(169). 10.3791/62045.

31. Fannin SW, et al. Aging Selectively Decreases Oxidative Capacity in Rat Heart Interfibrillar Mitochondria. Archives of Biochemistry and Biophysics. 1999;372(2):399–407.

32. Conrad R, Narayan K. Instance segmentation of mitochondria in electron microscopy images with a generalist deep learning model trained on a diverse dataset. cels. 2023;14(1):58–71.e5.

33. Belevich I, et al. Microscopy Image Browser: A Platform for Segmentation and Analysis of Multidimensional Datasets. PLoS Biol. 2016;14(1):e1002340.

34. Belevich I, Jokitalo E. DeepMIB: User-friendly and open-source software for training of deep learning network for biological image segmentation. PLoS Comput Biol. 2021;17(3):e1008374.

35. Adams RA, et al. Structural Analysis of Mitochondria in Cardiomyocytes: Insights into Bioenergetics and Membrane Remodeling. Curr Issues Mol Biol. 2023;45(7):6097–6115.

36. Riva A, et al. Structure of cristae in cardiac mitochondria of aged rat. Mech Ageing Dev. 2006;127(12):917–921.

37. Jiang Y-F, et al. Electron tomographic analysis reveals ultrastructural features of mitochondrial cristae architecture which reflect energetic state and aging. Sci Rep. 2017;7:45474.

38. Giacomello M, et al. The cell biology of mitochondrial membrane dynamics. Nat Rev Mol Cell Biol. 2020;21(4):204–224.

39. Davies KM, et al. Structure of the yeast F _1_ F _o_ -ATP synthase dimer and its role in shaping the mitochondrial cristae. Proc Natl Acad Sci USA. 2012;109(34):13602–13607.

40. Sebastián D, Palacín M, Zorzano A. Mitochondrial Dynamics: Coupling Mitochondrial Fitness with Healthy Aging. Trends in Molecular Medicine. 2017;23(3):201–215.

41. Hinton A, et al. Mitochondrial Structure and Function in Human Heart Failure. Circulation Research. 2024;135(2):372–396.

42. Chen L, et al. Mitochondrial OPA1, apoptosis, and heart failure. Cardiovascular Research. 2009;84(1):91–99.

43. Evidence of Glycolysis Up-Regulation and Pyruvate Mitochondrial Oxidation Mismatch During Mechanical Unloading of the Failing Human Heart: Implications for Cardiac Reloading and Conditioning. JACC: Basic to Translational Science. 2016;1(6):432–444.

44. Chaanine AH, et al. Mitochondrial Morphology, Dynamics, and Function in Human Pressure Overload or Ischemic Heart Disease With Preserved or Reduced Ejection Fraction. Circulation: Heart Failure. 2019;12(2):e005131.

45. Badolia R, et al. The Role of Nonglycolytic Glucose Metabolism in Myocardial Recovery Upon Mechanical Unloading and Circulatory Support in Chronic Heart Failure. Circulation. 2020;142(3):259–274.

46. Han YS, et al. Alterations in cardiac contractile and regulatory proteins contribute to age-related cardiac dysfunction in male rats. Physiol Rep. 2024;12(16):e70012.

47. Regmi SG, Rolland SG, Conradt B. Age-dependent changes in mitochondrial morphology and volume are not predictors of lifespan. Aging. 2014;6(2):118–130.

48. Amchenkova AA, et al. Coupling membranes as energy-transmitting cables. I. Filamentous mitochondria in fibroblasts and mitochondrial clusters in cardiomyocytes. J Cell Biol. 1988;107(2):481–495.

49. Glancy B, et al. Power Grid Protection of the Muscle Mitochondrial Reticulum. Cell Rep. 2017;19(3):487–496.

50. Eisner V, et al. Mitochondrial fusion dynamics is robust in the heart and depends on calcium oscillations and contractile activity. Proceedings of the National Academy of Sciences. 2017;114(5):E859–E868.

51. Dlasková A, et al. Mitochondrial cristae narrowing upon higher 2-oxoglutarate load. Biochimica et Biophysica Acta (BBA) - Bioenergetics. 2019;1860(8):659–678.

52. Varanita T, et al. The Opa1-Dependent Mitochondrial Cristae Remodeling Pathway Controls Atrophic, Apoptotic, and Ischemic Tissue Damage. Cell Metabolism. 2015;21(6):834–844.

53. Wolf DM, et al. Individual cristae within the same mitochondrion display different membrane potentials and are functionally independent. EMBO J. 2019;38(22):e101056.

54. Morris S, et al. Decreased ATP synthase activity is linked to altered spatiotemporal organisation of ATP Synthase in a cellular cardiomyocyte senescent model. 2024.

55. Song M, et al. Mitochondrial Fission and Fusion Factors Reciprocally Orchestrate Mitophagic Culling in Mouse Hearts and Cultured Fibroblasts. Cell Metabolism. 2015;21(2):273–286.

56. Song M, et al. Abrogating Mitochondrial Dynamics in Mouse Hearts Accelerates Mitochondrial Senescence. Cell Metabolism. 2017;26(6):872–883.e5.

57. Chen H, et al. Titration of mitochondrial fusion rescues *Mff* -deficient cardiomyopathy. Journal of Cell Biology. 2015;211(4):795–805.

58. Sabbah HN, et al. Abnormalities of Mitochondrial Dynamics in the Failing Heart: Normalization Following Long-Term Therapy with Elamipretide. Cardiovasc Drugs Ther. 2018;32(4):319–328.

59. Tezze C, et al. Age-Associated Loss of OPA1 in Muscle Impacts Muscle Mass, Metabolic Homeostasis, Systemic Inflammation, and Epithelial Senescence. Cell Metabolism. 2017;25(6):1374–1389.e6.

60. Cartes-Saavedra B, et al. OPA1 disease-causing mutants have domain-specific effects on mitochondrial ultrastructure and fusion. Proc Natl Acad Sci U S A. 2023;120(12):e2207471120.

61. Piquereau J, et al. Down-regulation of OPA1 alters mouse mitochondrial morphology, PTP function, and cardiac adaptation to pressure overload. Cardiovascular Research. 2012;94(3):408–417.

62. Chen L, et al. OPA1 Mutation and Late-Onset Cardiomyopathy: Mitochondrial Dysfunction and mtDNA Instability. JAHA. 2012;1(5):e003012.

63. Frezza C, et al. OPA1 Controls Apoptotic Cristae Remodeling Independently from Mitochondrial Fusion. Cell. 2006;126(1):177–189.

64. Afzal N, et al. Effect of crista morphology on mitochondrial ATP output: A computational study. Curr Res Physiol. 2021;4:163–176.

65. Baker N, Patel J, Khacho M. Linking mitochondrial dynamics, cristae remodeling and supercomplex formation: How mitochondrial structure can regulate bioenergetics. Mitochondrion. 2019;49:259–268.

66. Jang S, Javadov S. OPA1 regulates respiratory supercomplexes assembly: The role of mitochondrial swelling. Mitochondrion. 2020;51:30–39.

67. Gómez LA, et al. Supercomplexes of the mitochondrial electron transport chain decline in the aging rat heart. Archives of Biochemistry and Biophysics. 2009;490(1):30–35.

68. Divakaruni AS, Jastroch M. A practical guide for the analysis, standardization, and interpretation of oxygen consumption measurements. Nat Metab. 2022;4(8):978–994.

69. Ranjbarvaziri S, et al. Altered Cardiac Energetics and Mitochondrial Dysfunction in Hypertrophic Cardiomyopathy. Circulation. 2021;144(21):1714–1731.

70. Xue R-Q, et al. Regulation of mitochondrial cristae remodelling by acetylcholine alleviates palmitate-induced cardiomyocyte hypertrophy. Free Radical Biology and Medicine. 2019;145:103–117.

71. Bernardi P, et al. Identity, structure, and function of the mitochondrial permeability transition pore: controversies, consensus, recent advances, and future directions. Cell Death Differ. 2023;30(8):1869–1885.

72. Deerinck TJ, et al. High-performance serial block-face SEM of nonconductive biological samples enabled by focal gas injection-based charge compensation. J Microsc. 2018;270(2):142–149.

73. Sikkel MB, et al. Hierarchical statistical techniques are necessary to draw reliable conclusions from analysis of isolated cardiomyocyte studies. Cardiovascular Research. 2017;113(14):1743–1752.

