## Supplementary Material for "Multi-scale mitochondrial cristae remodeling links Opa1 downregulation to reduced OXPHOS capacity in aged hearts"

### SUPPLEMENTAL FIGURES

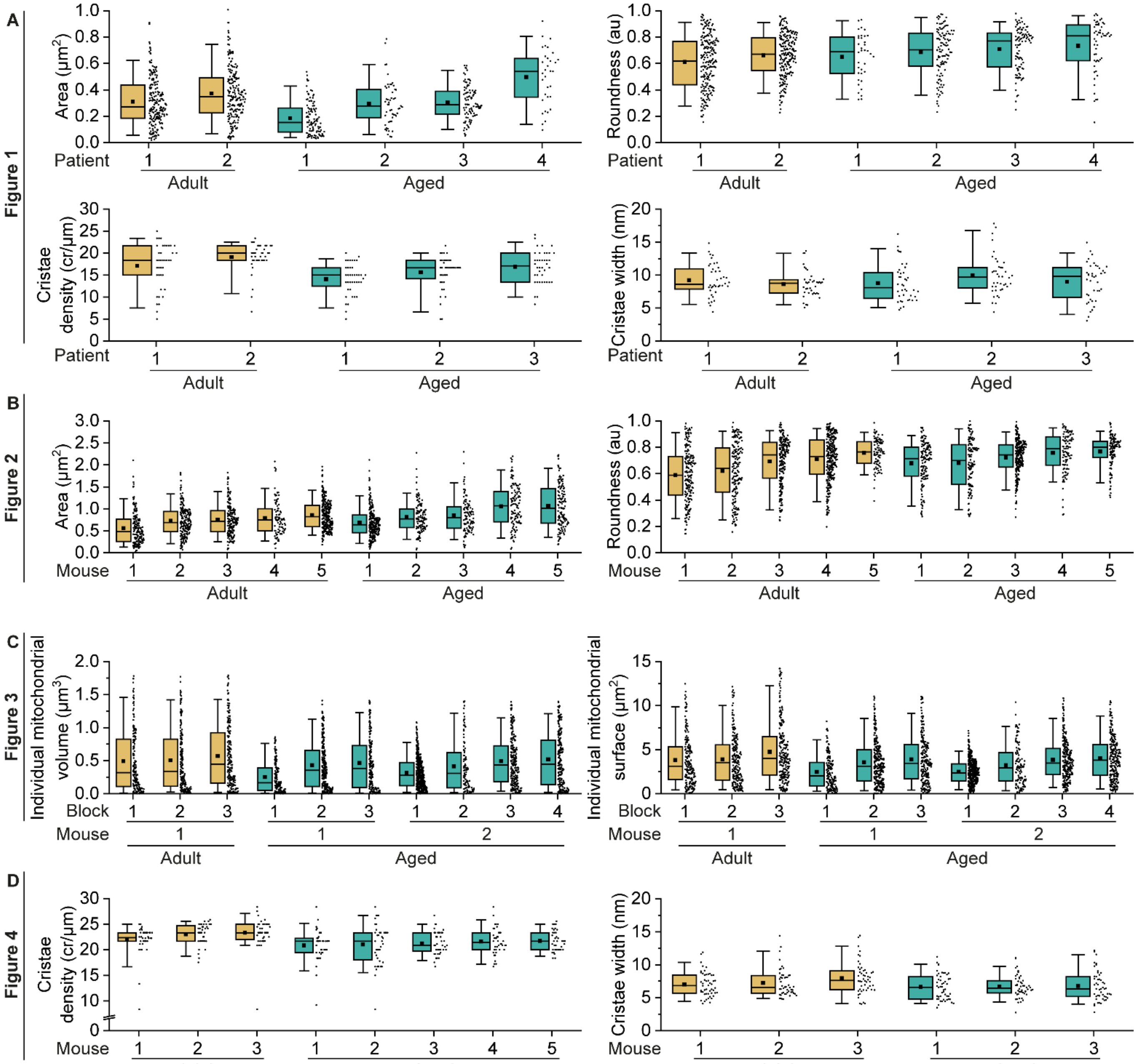


**Supplemental Figure 1. Hierarchical data distribution across different figures. (A)** Data distribution from patients of **Figure 1**. **(B)** Data distribution from mice of **Figure 2**. **(C)** Data distribution from mice and their respective tissue blocks from **Figure 3**. **(D)** Data distribution from mice of **Figure 4**. Box: P25 and P75. Whiskers: P5 and P95. Line: Median. Square: Mean. Each datapoint represents a mitochondrion.

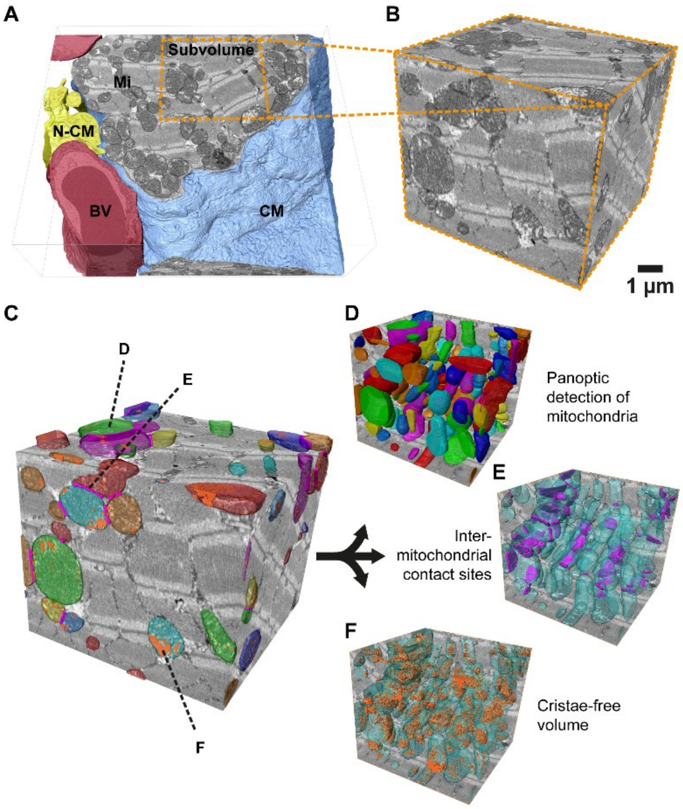


**Supplemental Figure 2. Sub-volume analysis of IMF by SBF-SEM. (A)** Volumetric representation of the reconstructed SBF-SEM data containing the cardiomyocyte and parts of the cardiac environment. Blood vessel (BV), extracellular immune components and fibroblasts (N-CM), cardiomyocyte (CM), and mitochondria (Mi). Dimensions = 20×20×6 $\mu m^3$. **(B)** Enlarged 3D orthoview of representative sub-volume used for subsequent measurements, indicated by orange inset in **(A)**. Targeted region contains IMF. Dimensions = 9.5×9.5×6 $\mu m^3$. **(C)** Identical volume as in **(B)** with annotated analysis targets: panoptically segmented, multi-colored mitochondria used for volume and surface area measurements **(D)**; inter-mitochondrial contact sites (IMCS) areas in magenta **(E)**; Cristae-free volume in orange **(F)**. **(D-F)** 3D surface reconstructions of respective segmentations highlighted in **(C)**.

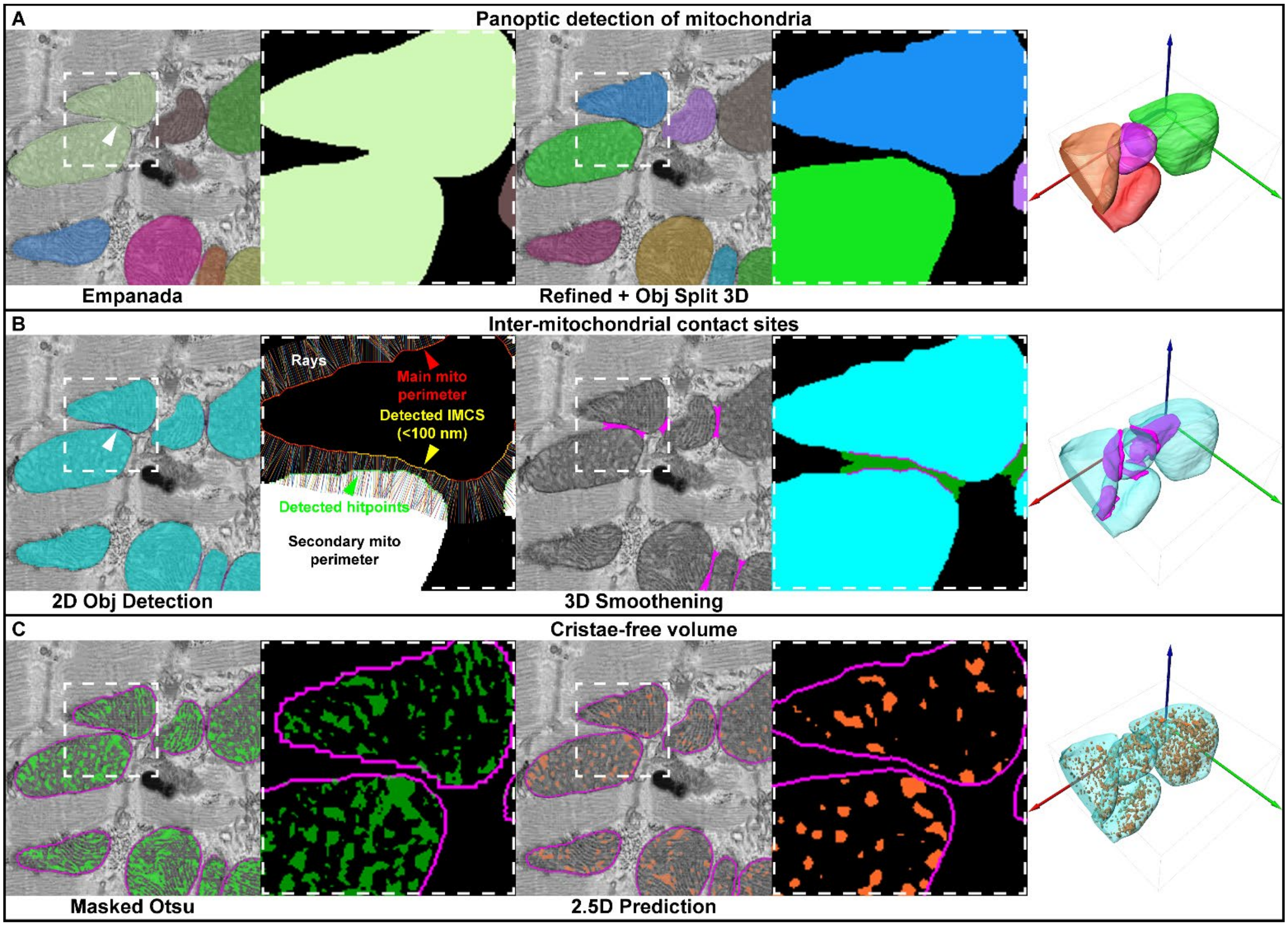


**Supplemental Figure 3. Segmentation workflow of SBF-SEM data to study mitochondrial ultrastructure within cardiomyocytes. (A)** Panoptic segmentation prediction of mitochondria via Empanada (left); 3D object separation in MIB (center); 3D reconstruction of the corresponding volume (right). Arrowheads highlight a case of oversegmentation. **(B)** MIB MCcalc-Plugin automatic detection of IMCS for each slice in 2D (left); IMCS smoothening based on the detected 2D juxta-membranes; 3D reconstruction of the corresponding volume (right). Arrowheads highlight a detected IMCS. **(C)** Otsu-based binarization of masked areas to detect cristae-free volume (green) within mitochondria (magenta) (left); 2.5D deep learning model prediction to consider three-dimensional morphology of mitochondrial matrix more independent of global contrast differences (orange) (center); 3D reconstruction of the corresponding volume (right). White boxes indicate an inset for a detailed view of the underlying binary model masks.

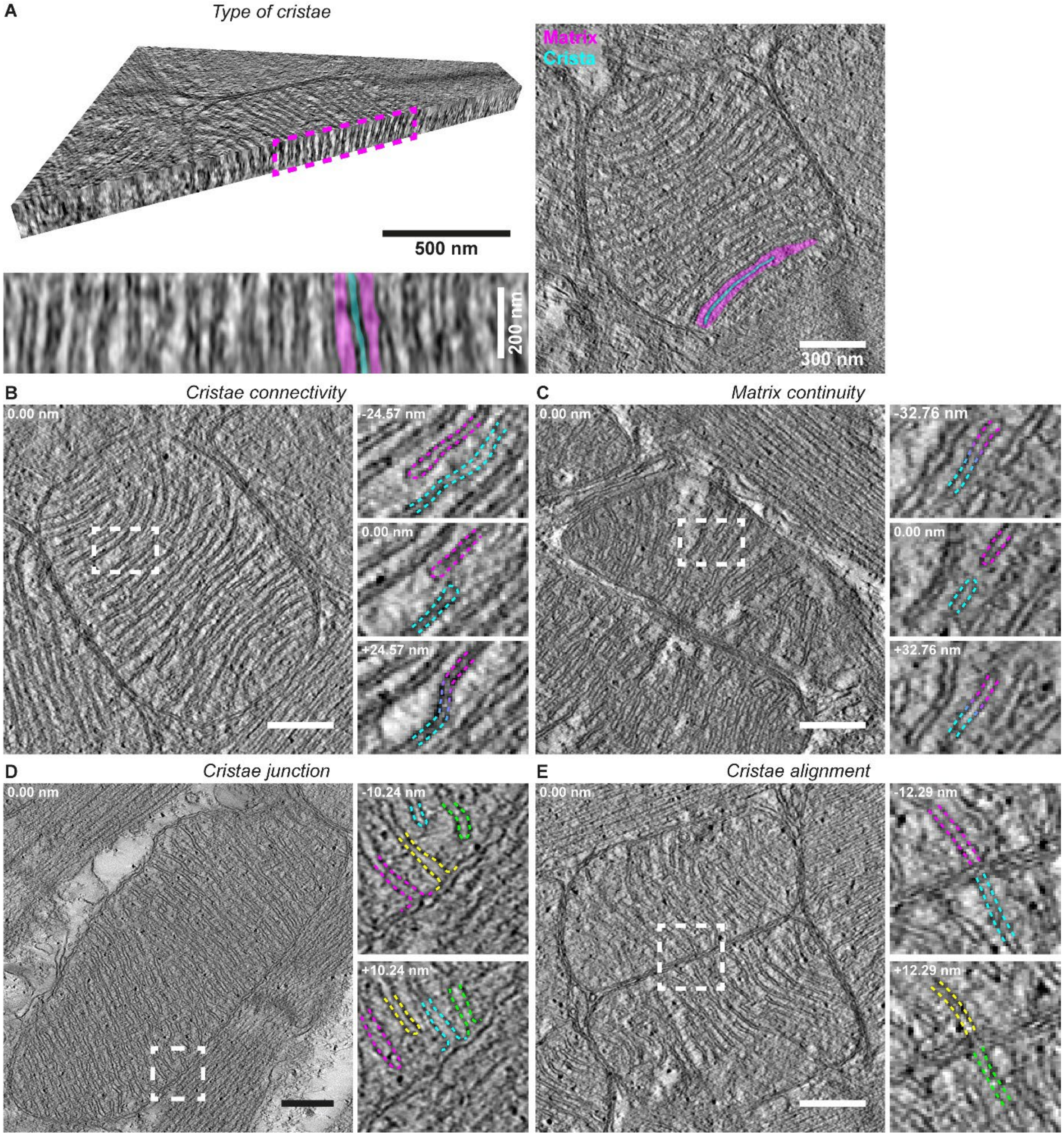


**Supplemental Figure 4. Electron tomography applications for mitochondrial ultrastructure studies.** Methods to identify type of cristae **(A)**, cristae connectivity **(B)**, matrix connectivity **(C)**, cristae junction **(D)**, and cristae alignment **(E)**. Tomograms were acquired from ~200 nm sections at 28000X with an XYZ resolution of 4.10 nm **(A-C, E)**, and 2.05 nm **(D)**.

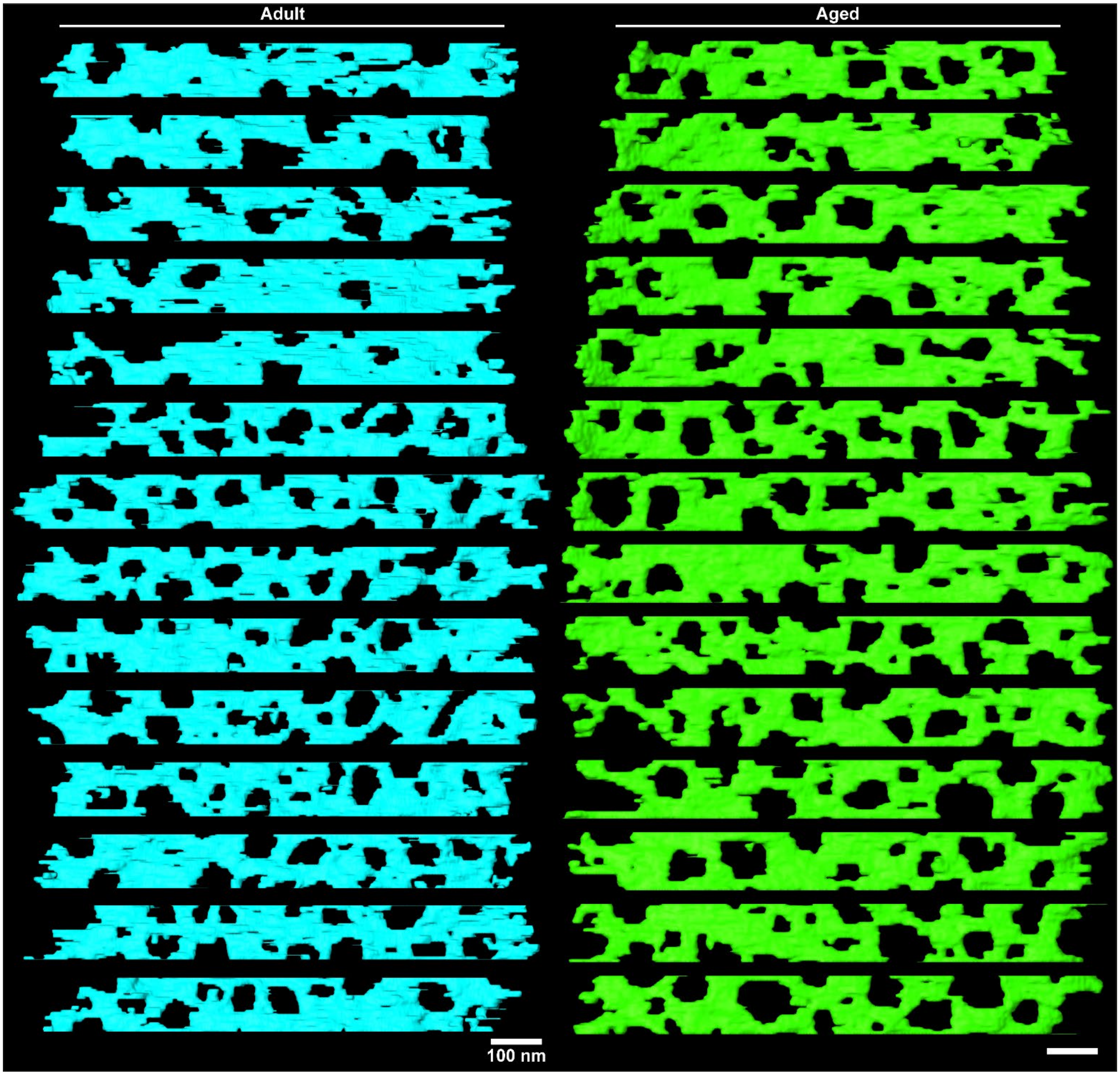


**Supplemental Figure 5. Mitochondrial lamellar cristae from mice hearts exhibit an increase in fenestration size with aging.** Mitochondrial lamellar cristae from adult and aged mice hearts displayed from a frontal view. Cristae fenestrations can be observed as interruptions of the lamellar cristae varying in size.

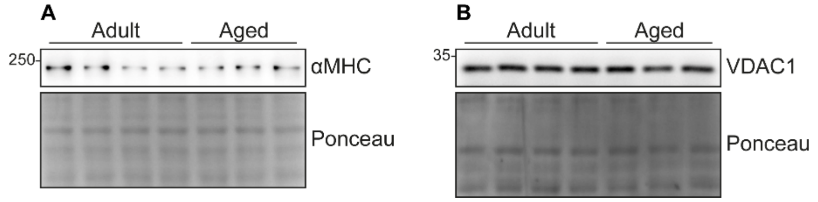


**Supplemental Figure 6. Mitochondrial mass remains constant in mice aged heart. (A)** Western blot of αMHC normalized against Ponceau staining as loading control. **(B)** Western blot of VDAC1 normalized against Ponceau staining as loading control.

### SUPPLEMENTAL VIDEOS

**Supplemental Video 1. Segmentation and analysis pipeline of an adult mice heart sample**. SBF-SEM volumes focusing on the IFM were segmented using Empanada and a semi-automatic curation approach with the 3D object splitting tool in MIB. The segmentation data was then used to calculate volume and surface area, represented in the video as mitochondria with multiple colors. For IMCS, the MIB MCcal plugin was used to analyze contact points and distance distribution, denoted as magenta sheets. Cristae-free volume was assessed using the curated segmentation with a selective threshold, represented as three-dimensional golden dots.

**Supplemental Video 2. Segmentation and analysis pipeline of an aged mice heart sample.** As in **Supplemental Video 1**, IFM were segmented using Empanada and curated with MIB. The segmentation data enabled volume and surface area calculations, shown as multicolored mitochondria. IMCS analysis were performed with MCcal plugin, denoted as magenta sheets. Cristae-free volume was calculated by applying a threshold was represented as 3D golden dots.

**Supplemental Video 3. Tomographic volume and 3D segmentation of mitochondrion from an adult mice heart sample.** A three-dimensional reconstruction of a mitochondrion was generated through manual segmentation using the software Amira. The video highlights the detailed structure, including lamellar cristae and cristae junctions, providing insights into mitochondrial organization.

**Supplemental Video 4. Tomographic volume and 3D segmentation of mitochondrion from an aged mice heart sample.** As in **Supplemental Video 3**, the video shows a three-dimensional reconstruction of mitochondrion, showing lamellar cristae and cristae junctions.

**SUPPLEMENTAL TABLES**

**Supplemental Table 1.** Antibody detailed information.

| Antibody | Manufacturer | Catalog number | Dilution |
|---|---|---|---|
| Mfn1 | Donated by Dr. Richard Youle | - | 1:5000 |
| Mfn2 | Abcam | 50838 | 1:2000 |
| Opa1 | BD Biosciences | 612607 | 1:1000 |
| Drp1 | BD Biosciences | 611113 | 1:2000 |
| pDrp1 S616 | Cell Signaling | 3455S | 1:2000 |
| pDrp1 S637 | Cell Signaling | 4867S | 1:2000 |
| Fis1 | Proteintech | 10956-1-AP | 1:1000 |
| Mff | Proteintech | 17090-1-AP | 1:2000 |
| Mid49 | Proteintech | 16413-1-AP | 1:1000 |
| Mid51 | Proteintech | 20164-1-AP | 1:1000 |
| Mic60 | Proteintech | 10179-1-AP | 1:1000 |
| Mic19 | Proteintech | 25625-1-AP | 1:1000 |
| Mic10 | Abcam | 84969 | 1:1000 |
| PGC1-α | Novus | NBP1-04676 | 1:1000 |
| OXPHOS cocktail | Abcam | 110413 | 1:1000 |
| ATP5B | Abcam | 170947 | 1:5000 |
| ATP5I | Abcam | 122241 | 1:5000 |
| VDAC1 | Cell Signaling | D73D12 | 1:1000 |
| GAPDH | Proteintech | 60004-1 | 1:2000 |
| αMHC | Hybridoma Bank | MF20 | 1:5 |
| Anti-Mouse | Jackson ImmunoResearch | 115035068 | 1:5000 |
| Anti-Rabbit | Jackson ImmunoResearch | 111035045 | 1:5000 |

**Supplemental Table 2.** Primer sequence information.

| Gene | Species | Primer Forward | Primer Reverse |
|---|---|---|---|
| *Mfn1* | Mouse | 5'-CCA CAA GCT GTG TTC GGA TTT-3' | 5'-CAC CCT CTG TGC ATT TGT GG-3' |
| *Mfn2* | Mouse | 5'-TGT GCT GAC TTT CAG GAG GAC-3' | 5'-TGC AGC CTA GCA AGG CCC G-3' |
| *Opa1* | Mouse | 5'-CAG GAG AAG TAG ACT GTG TC-3' | 5'-TGT GAC TTT ATT TTG CAC GG-3' |
| *Mic60* | Mouse | 5'-TCT GAC CTA GCT GGC AAA CTC-3' | 5'-ACT GCA GAG TCA AAG GTC CG-3' |
| *Mic10* | Mouse | 5'-TTT GGT TCT GGC GTG GGA TT-3' | 5'-TGG CAC CAT CAC AGG ATG TC-3' |
| *Drp1* | Mouse | 5'-GCT ATG GTG AAC CGG TGG AT-3' | 5'-TGC GGT TCC TTC AAT CGT GT-3' |
| *Fis1* | Mouse | 5'-AAG TAT GTG CGA GGG CTG TT-3' | 5'-TGG CCT TAT CAA TCA GGC GTT-3' |
| *Mff* | Mouse | 5'-TGG AGC TAA TCT TTC CTC TGC C-3' | 5'-CAA GAT CTG CTG GTA AGC CCT A-3' |
| *GAPDH* | Mouse | 5'-AGG TCG GTG TGA ACG GAT TTG-3' | 5'-TGT AGA CCA TGT AGT TGA GGT CA-3' |
| *MFN1* | Human | 5'-GTT GGA GCG GAG ACT TAG CA-3' | 5'-GCA GTA ATC GCC TTC TTA GCC-3' |
| *OPA1* | Human | 5'-CTG GTA AAC GCG TTC AAC-3' | 5'-GAG CTG ATT ATG AGT ACG ATT TTA ATT-3' |
| *DRP1* | Human | 5'-GTC ATG GAG GCG CTA ATT CC-3' | 5'-TCC CAC TAC GAC GAT TTG AGG-3' |
| *MFF* | Human | 5'-GGG GAC AAA AGT GGC TCT CA-3' | 5'-GGC ACT ATC TGC TTC TGT GC-3' |
| *MIC60* | Human | 5'-TGG AGC TGG CCT TTT GTT TG -3' | 5'- TGG TTT TCT CTA CAC TTT CCC G-3' |
| *18S* | Human | 5'-GCA GAA TCC ACG CCA GTA CAA G-3' | 5'-GCT TGT TGT CCA GAC CAT TGG C-3' |
